# The Evolution of SlyA/RovA Transcription Factors From Repressors to Counter-Silencers in *Enterobacteriaceae*

**DOI:** 10.1101/369546

**Authors:** W. Ryan Will, Peter Brzovic, Isolde Le Trong, Ronald E. Stenkamp, Matthew B. Lawrenz, Joyce E. Karlinsey, William W. Navarre, Kara Main-Hester, Virginia L. Miller, Stephen J. Libby, Ferric C. Fang

## Abstract

Gene duplication and subsequent evolutionary divergence have allowed conserved proteins to develop unique roles. The MarR family of transcription factors (TFs) has undergone extensive duplication and diversification in bacteria, where they act as environmentally-responsive repressors of genes encoding efflux pumps that confer resistance to xenobiotics, including many antimicrobial agents. We have performed structural, functional, and genetic analyses of representative members of the SlyA/RovA lineage of MarR TFs, which retain some ancestral functions, including repression of their own expression and that of divergently-transcribed multidrug efflux pumps, as well as allosteric inhibition by aromatic carboxylate compounds. However, SlyA and RovA have acquired the ability to counter-silence horizontally-acquired genes, which has greatly facilitated the evolution of *Enterobacteriaceae* by horizontal gene transfer. SlyA/RovA TFs in different species have independently evolved novel regulatory circuits to provide the enhanced levels of expression required for their new role. Moreover, in contrast to MarR, SlyA is not responsive to copper. These observations demonstrate the ability of TFs to acquire new functions as a result of evolutionary divergence of both *cis*-regulatory sequences and *in trans* interactions with modulatory ligands.

## Introduction

As organisms adapt to new or changing environments, their regulatory networks must evolve to ensure that individual genes are appropriately expressed in response to environmental signals (1). An important mechanism for the evolution of conserved essential proteins, including transcription factors (TFs), is gene duplication, which allows the subsequent diversification of gene and protein families and the development of new functions (2). More than 50% of bacterial genes are believed to have descended from original duplication events (3-5), providing a broad foundation from which bacteria can evolve complex and adaptive traits.

The MarR family is an ancient family of TFs, predating the divergence of archaea and bacteria (6). It has undergone extensive gene duplication events, with recent estimates suggesting that bacteria encode an average of seven MarR TFs per genome (7). MarR TFs typically function as environmentally-responsive repressors of genes encoding efflux pumps that export xenobiotics, including many antimicrobial agents, and are defined by the presence of a winged helix-turn-helix (wHTH) DNA-binding domain (8). The prototypical MarR protein of *Escherichia coli* represses a single operon, *marRAB*, which encodes a transcriptional activator (MarA) required for the expression of the AcrAB efflux pump, which in turn confers resistance to β-lactams, quinolones, and tetracyclines (9-11). MarR is allosterically regulated by many small molecules, in particular small aromatic carboxylate compounds such as salicylate, which induce a structural change that reduces the affinity of MarR for DNA (10, 12-14) and derepresses the expression of its cognate promoters. A recent study suggests that MarR can also be inhibited by intracellular copper (Cu^2+^), which oxidizes a conserved cysteine residue at position 80, promoting the formation of disulfide bonds between MarR dimers and causing individual dimers to dissociate from DNA (15). Free copper is thought to be liberated from membrane-bound cytoplasmic proteins during envelope stress induced by antimicrobial agents. Dimerization of MarR TFs is required for DNA binding, as it allows these proteins to recognize palindromic sequences via the α4 recognition helix, which makes sequence-specific contacts with the major groove, while the wing makes sequence-independent contacts via the minor groove (16).

A duplication event producing the SlyA lineage of MarR TFs most likely resulted from an ancient horizontal gene transfer event or from intragenomic recombination of a MarR family TF prior to the divergence of the *Enterobactericeae*. SlyA has been best characterized in *Salmonella enterica* serovar Typhimurium, where it serves primarily to upregulate virulence genes (17-19). Although this contrasts with the classical repressive role of MarR TFs, work in our and other laboratories has demonstrated that SlyA positively regulates genes by a counter-silencing mechanism, in which repression of AT-rich promoters by the histone-like nucleoid-associated protein H-NS is relieved by SlyA (17, 20). SlyA cooperatively remodels the H-NS-DNA complex in concert with the response regulator PhoP (20, 21), which is activated by conditions found within phagosomal compartments, including low Mg^2+^ (22), acidic pH (23), and cationic antimicrobial peptides (24). SlyA orthologs, represented by Hor, Rap, and RovA in *Pectobacterium* (*Erwinia*), *Serratia*, and *Yersinia,* respectively (25), are conserved in nearly every species of *Enterobacteriaceae*, even including endosymbionts such as *Sodalis glossinidius* (26), which have undergone extensive gene loss and degenerative evolution (27). This high degree of conservation suggests that the SlyA lineage occupies an essential role in the regulatory network organization of *Enterobacteriaceae*. Although conclusive mechanistic evidence to demonstrate that other SlyA orthologs function as counter-silencers has not yet been obtained, existing evidence is strongly suggestive of counter-silencing, as several are known to up-regulate horizontally-acquired traits, which are generally repressed by H-NS, in a number of species, including *Yersinia* spp. (28-30), *Dickeya dadantii* (31), *Pectobacterium carotovorum* (25) and *Shigella flexneri* (32).

TFs can evolve in two ways: *in cis,* through their promoters and associated regulatory elements, both transcriptional and post-transcriptional, altering expression patterns to respond to different environmental and physiological stimuli, and *in trans*, affecting their interactions with cognate binding sites, other proteins, and regulatory ligands. We sought to understand the evolutionary transition of the SlyA/RovA TF lineage from the ancestral function of MarR family TFs as environmentally-responsive and dedicated repressors of small regulons to counter-silencers of extensive networks of horizontally-acquired genes, with a particular focus on *in cis* changes in gene expression and *in trans* changes in modulation by inhibitory ligands. Structural and comparative analyses of representative members of the SlyA lineage were performed to identify the evolutionary changes that allowed SlyA to adopt its new role. Here we show that SlyA has retained an ability to undergo conformational changes in response to aromatic carboxylates, regulate gene expression in an environmentally-responsive manner, and repress the expression of a linked drug efflux system. However, SlyA/RovA lineage genes have undergone extensive evolution *in cis* to support the higher levels of expression that are required for counter-silencing. Finally, we show that linked efflux pumps are not conserved in some *Enterobacteriaceae,* even though SlyA/RovA TFs have been evolutionarily retained, suggesting that these regulators have been conserved not due to their primordial role in regulating antimicrobial resistance but rather as a consequence of their counter-silencing function, which is essential to maintain the regulated expression of horizontally-acquired genes in *Enterobacteriaceae*.

## Results

### Salicylate-mediated inhibition of SlyA activity

As environmentally-responsive repressors whose conformation and regulatory actions are modulated by small aromatic carboxylates (12, 14), MarR family TFs are inhibited by salicylate *in vitro* (12). In their structural analyses of SlyA-DNA interactions, Dolan *et al*, (33) inferred from our structural data (see below) that salicylate might regulate SlyA. Using electrophoretic mobility shift assays, they demonstrated that salicylate inhibits DNA-binding by SlyA. To confirm that this influences the function of SlyA as a transcriptional regulator, we performed *in vitro* transcription assays (IVTs) of *slyA* and the divergently transcribed *ydhIJK* efflux pump operon. Supercoiled plasmid DNA containing the *slyA-ydhIJK* region was incubated with RNA polymerase (RNAP) and increasing SlyA concentrations in the presence or absence of salicylate. SlyA repressed *slyA* transcription approximately 5.3-fold, while *ydhI* transcription was inhibited ∼19-fold (Figure 1A, B). The addition of 2mM sodium salicylate reduced SlyA-mediated repression to 2.8-fold and 3.2-fold, respectively, indicating that the sensitivity to aromatic carboxylates observed in classical MarR TFs has been retained by SlyA.

**Figure 1.**
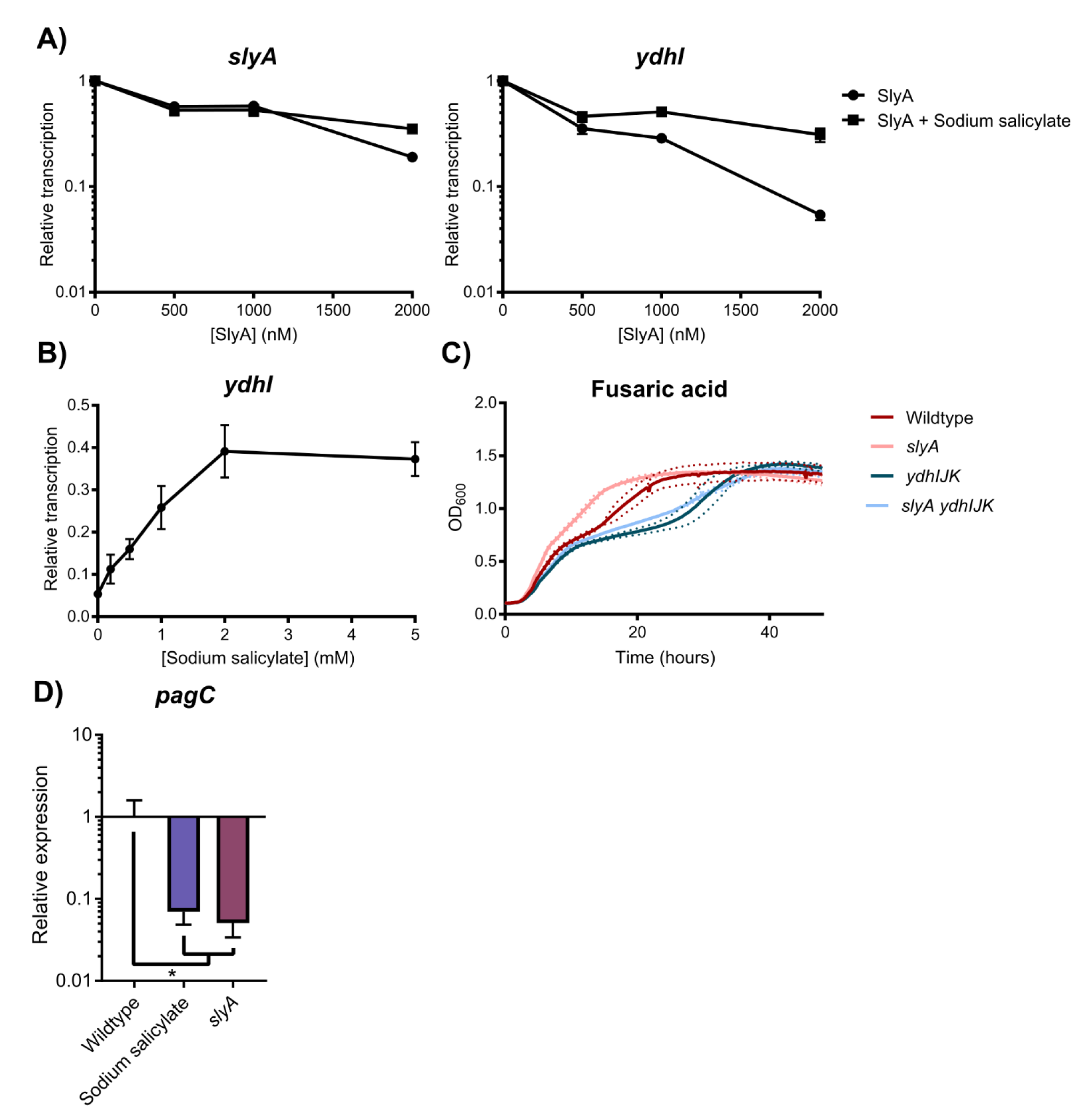
SlyA retains the ancestral functions of MarR family TFs. To determine whether SlyA is an autorepressor like other members of the MarR family, *in vitro* transcription (IVT) assays were performed on supercoiled template DNA containing *slyA* and the divergently expressed *ydhI* gene in the presence of increasing SlyA concentrations (A). Reactions were performed with or without 2mM sodium salicylate to determine whether SlyA is inhibited by small aromatic carboxylate compounds like other MarR TFs. (IVT analysis of SlyA-mediated regulation of *pagC* is shown in Fig. S1.) To determine whether SlyA exhibits a dose-dependent response to salicylate, IVT assays were performed on *ydhI* in the presence of 2µM SlyA and increasing concentrations of sodium salicylate (B). Wildtype, *slyA, ydhIJK*, and *slyA ydhIJK* cultures were grown in the presence of 30 µg/ml fusaric acid and cell density (OD_600_) measured over time to determine if *ydhIJK* encodes a functional anti-microbial efflux pump (C). Data represent the mean (solid line) of three independent experiments, each consisting of three replicates. Dashed lines represent the SD. (Growth curves in the absence of fusaric acid is shown in Fig. S2.) To determine whether salicylate also inhibits SlyA counter-silencing activity, *pagC* transcripts were quantified by qRT-PCR from early stationary phase (OD_600_≈2.0) cultures in minimal N-medium containing 10 µM MgCl_2_ in the presence or absence of 2 mM sodium salicylate (D). Transcript levels are normalized to *rpoD*, and data represent the mean ± SD; n≥3. Asterisk indicates P=0.05.

We then confirmed that the *ydhIJK* operon encodes a functional antimicrobial efflux pump by growing wildtype, *slyA, ydhIJK*, and *slyA ydhIJK* mutant strains in the presence of an aromatic carboxylate with antimicrobial activity, fusaric acid (Figure 1C). Both the *ydhIJK* and *slyA ydhIJK* mutant strains exhibited delayed growth in the presence of fusaric acid relative to the wildtype strain, suggesting that *ydhIJK* confers fusaric acid resistance in wildtype *S*. Typhimurium. Conversely, *slyA* mutants exhibit improved growth in the presence of fusaric acid, as predicted when *ydhIJK* is derepressed. Collectively, these observations indicate that the *S.* Typhimurium SlyA TF has retained ancestral functions characteristic of the MarR family.

To determine whether salicylate inhibits SlyA-mediated counter-silencing as well as repression, expression of the counter-silenced *pagC* gene was measured by qRT-PCR in the presence or absence of 2mM sodium salicylate (Figure 1D). Wildtype cultures grown in salicylate phenocopied a *slyA* mutant strain, with a >14-fold reduction in *pagC* expression, indicating that salicylate is a general allosteric inhibitor of SlyA function, most likely inducing a structural change that reduces affinity for DNA as described in other MarR TFs (8, 10). Unexpectedly, SlyA retained an ability to interact with DNA upstream of the *ydhI* promoter even in the presence of salicylate (Figure 2 - figure supplement 1), but this interaction may represent non-specific interactions with the wing domain, as salicylate inhibited SlyA interaction with a 12 bp DNA region that is highly homologous (75% identity) to the consensus high affinity binding site (19, 34, 35), centered near the −35 of the *ydhI* TSS.

**Figure 2.**
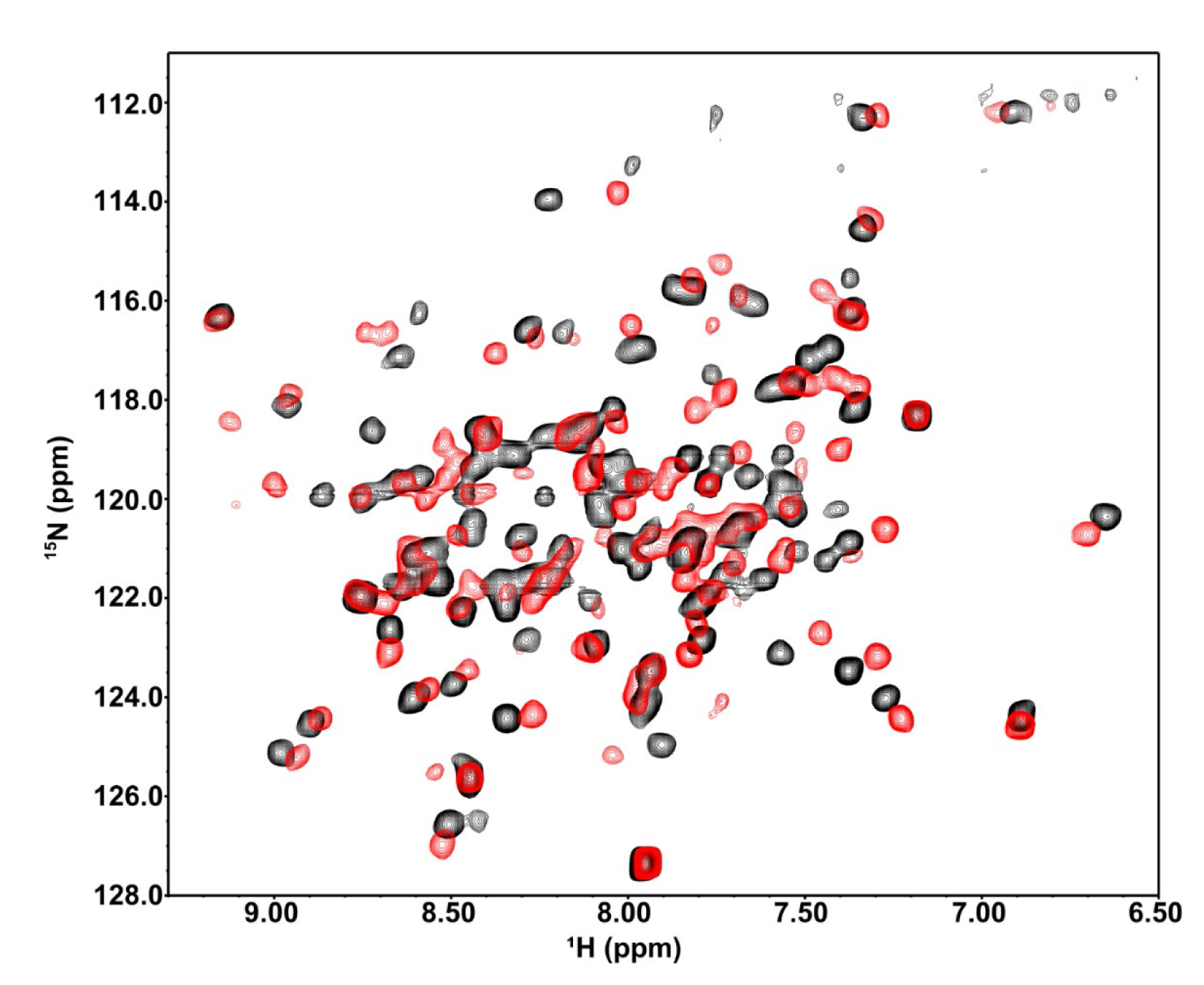
SlyA undergoes significant structural alterations upon salicylate binding. To assess salicylate-induced SlyA structural changes, ^1^H,^15^N-TROSY NMR spectroscopy was performed on 0.3mM uniformly-labelled ^15^N-SlyA in the absence (black) or presence (red) of 2mM sodium salicylate. (Differential DNA Footprinting Analysis, or DDFA, of the SlyA-DNA interaction in the presence or absence of salicylate is shown in Figure 2 – figure supplement 1 Ligand binding analysis is shown in Figure 2 – figure supplement 2.)

To determine whether salicylate-mediated inhibition correlates with structural changes in SlyA, ^1^H,^15^N-TROSY NMR spectra of *S.* Typhimurium SlyA in the presence or absence of salicylate were collected. The apo-SlyA spectrum is well dispersed (Figure 2), with ∼85% of the expected resonances observed. However, wide variation among individual resonances with respect to peak width and intensity may signify [1] weak non-specific interactions between SlyA dimers, [2] varying rates of exchange of amide protons with solvent, or [3] conformational exchange. Previous studies of MarR found that apo-MarR is also highly disordered (8), suggesting that the observed spectral characteristics of apo-SlyA can be ascribed to the presence of multiple conformational states in solution. Addition of salicylate to SlyA induces chemical shift perturbations throughout the spectrum and nearly 90% of expected resonances are now observed. The large-scale changes in chemical shifts show that backbone amides throughout the protein are stabilized in a different environment relative to apo-SlyA. These observations are consistent with ligand binding to SlyA dimers inducing global structural changes, likely stabilizing a single protein conformation in solution, a conformation that is no longer able to interact with specific high-affinity DNA binding sites. The affinity of the SlyA-ligand interaction was determined via the quenching of intrinsic tryptophan fluorescence in the presence of increasing concentrations of benzoate, an analogous small aromatic carboxylate compound (Figure 2 - figure supplement 2). Benzoate induces similar chemical shift perturbations to those observed in response to salicylate but does not cause inner filter effects that interfere with fluorescence measurement, as there is relatively little overlap between the UV spectra of benzoate and SlyA, in contrast to salicylate. Although this assay was not able to differentiate between multiple SlyA-benzoate interactions, the K_D_ of the SlyA-benzoate interaction was determined to be ∼40µM, which is similar to the previously determined affinities for EmrR/MprA and HucR (1-10µM), and significantly stronger than that of the MarR-salicylate interaction (0.5-1mM) (13, 36).

### Crystal structure of salicylate-SlyA

To further analyze the mechanism of allosteric inhibition of SlyA, the structure of the salicylate-SlyA co-crystal was determined. Studies by other groups have previously determined the structure of apo-SlyA and the SlyA bound to DNA (33), demonstrating that SlyA is similar in overall structure to other MarR proteins, consisting of six alpha helices (Figure 3A). Helices α1, α5, and α6 make up the dimerization domain, while α3 and α4, along with the wing region between α4 and α5 comprise the wHTH DNA-binding domain. These two domains are separated by α2. Dolan *et al.* (33) previously observed that the two recognition helices of the apo-SlyA dimer are only ∼15Å apart, in a closed conformation. During interaction with a high-affinity binding site, the helices move a significant distance to accommodate the 32Å distance between major grooves.

**Figure 3.**
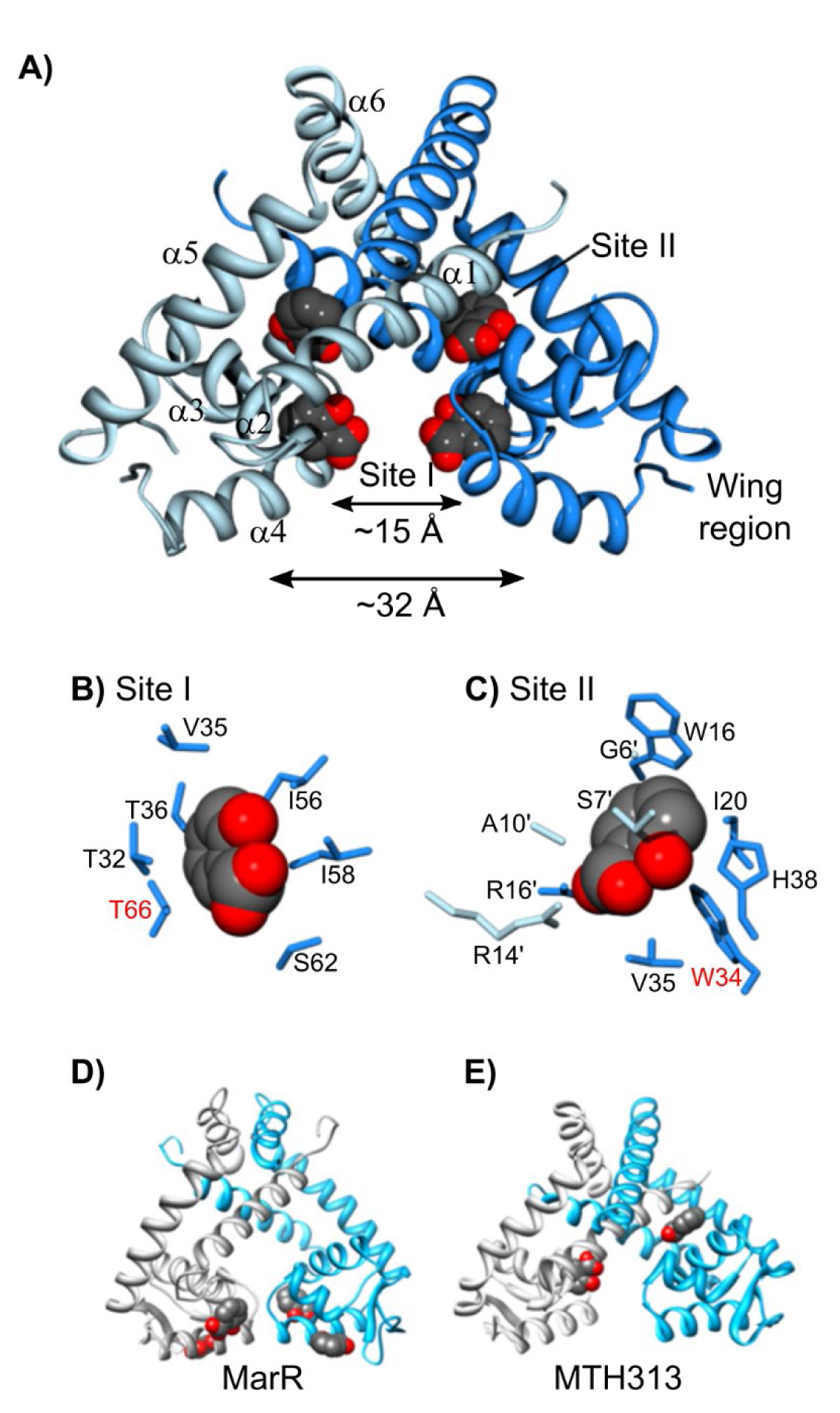
Structure of the SlyA dimer bound to salicylate. The crystal structure of the SlyA dimer with salicylate bound at sites I and II was determined at a resolution of 2.3Å (A). Residues involved in coordinating bound salicylates are shown for sites I (B) and II (C) Residues required for salicylate-mediated inhibition of SlyA (see Fig. 4) are highlighted in red. The structures of salicylate-bound MarR (D; PDB entry 1JGS) and MTH313 (E; PDB entry 3BPX) dimers are shown for comparison. (An image of the SlyA-salicylate crystal is shown in Figure 3 - figure supplement 1.)

SlyA formed large crystals in the presence of 75mM sodium salicylate (Figure 3 - figure supplement 1). However, we were unable to obtain usable crystals of apo-SlyA. Diffraction data set and refinement statistics are summarized in Tables S1 and S2, respectively. The two SlyA molecules in the asymmetric unit form two different SlyA dimers in this crystal form. Space group symmetry operations generate the other subunit in each dimer. The dimers are very similar in structure, and further discussion will focus on the dimer formed by polypeptide chain A. We were unable to observe electron density for the tips of the wings, so these regions are absent from our structural model. Difference electron density maps (|Fo|-|Fc|) identified two salicylate molecules bound per SlyA monomer at sites referred to as Site I and Site II (Figure 3A-C). Salicylate molecules interact with these sites via hydrophobic interactions with their aromatic rings, while the carboxylate and hydroxyl groups are positioned to interact with the solvent. Site I is composed of residues from α2, α3, and α4, as well as I58 in the loop between α3 and α4, and is well positioned to sterically inhibit DNA binding. It should be noted that the residue numbers in the deposited PDB file are not in register with the residue numbers in this text which are based on alignment of SlyA orthologs. Comparison with apo-SlyA and SlyA-DNA structures (33) indicates that this salicylate molecule causes the α4 recognition helix to rotate by ∼35° around its axis, disrupting specific contacts with the DNA major groove. Site II is formed by residues from both dimer subunits, almost completely sequestering the salicylate molecule from the solvent. The buried polar groups of salicylate interact with S7’ and R14’ in α1 of one subunit and R17 in α1 and H38 in α2 from the other subunit. A third salicylate binding site was observed on the surface of each subunit of the dimer. However, this site is adjacent to SlyA residues involved in crystal packing contacts and may not be biologically relevant.

### Mutational analysis of allosteric inhibition of SlyA

We constructed a series of *slyA* alleles with site-specific mutations of the salicylate-binding pocket in order to test the functional significance of salicylate interactions *in vivo*. When wildtype *slyA* is expressed *in trans* from its native promoter, *pagC* expression increases over 350-fold in inducing medium containing 10µM MgCl_2_ compared to cultures grown with salicylate (Figure 4A). We tested 8 different mutant alleles for changes in counter-silencing activity in response to salicylate. One allele substituting an alanine for tyrosine 66 in site I (T66A), resulted in complete abrogation of salicylate-mediated inhibition, suggesting that T66 is essential for salicylate binding. However the T66A mutation also decreased *pagC* expression in the absence of salicylate over 25-fold, indicating that it is required for the wildtype activity of SlyA. A second mutant, W34A in site II, also exhibited reduced salicylate-mediated repression. The analysis of these mutants indicates that both site I and site II bind salicylate *in vivo*, and that both sites influence SlyA activity. Notably, both T66 and W34 are absolutely conserved in over 55 enterobacterial genera examined in this study (see below), suggesting that these residues are important for SlyA function throughout the *Enterobacteriaceae* (Table S3).

**Figure 4.**
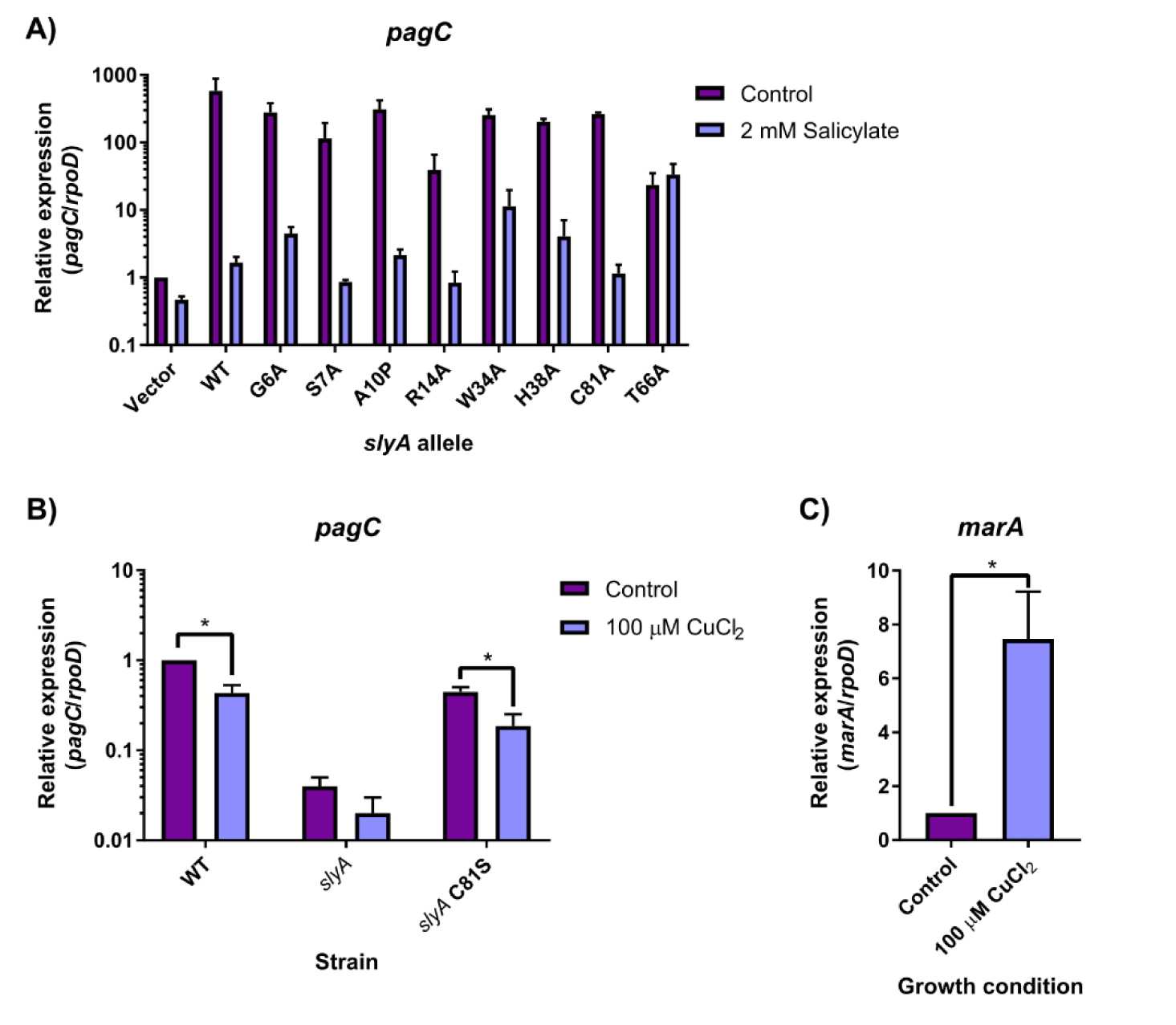
Genetic analyses of allosteric inhibition of SlyA. Selected residues involved in coordinating SlyA-bound salicylate were mutated in pSL2143, a plasmid expressing *slyA* from its native promoter, and assayed for their ability to restore counter-silencing in a chromosomal *slyA* mutant. Cultures were induced in 10µM MgCl_2_-containing medium in the presence or absence of 2mM sodium salicylate (A). Levels of *pagC* mRNA were quantified by qRT-PCR and normalized to *rpoD*. The mutant strain carrying the empty pWSK29 vector was included as a control. To determine if SlyA was directly inhibited by Cu^2+^ via C81, *pagC* expression was quantified in wild-type or isogenic *slyA* and *slyA* C81S mutant *S*. Typhimurium strains in inducing medium in the presence or absence of 100µM CuCl_2_ (B). To confirm that Cu^2+^-mediated inhibition occurs under these conditions, *marA* transcription was also quantified in the presence or absence of 100µM CuCl_2_. Data represent the mean ± SD; n=3. Asterisks indicate P < 0.005.

To determine whether salicylate directly inhibits SlyA *in vivo* or stimulates the release of intracellular Cu^2+^ from membrane bound proteins to promote disulfide bond formation, we mutated the single cysteine residue in SlyA, C81. SlyA C81A exhibited a similar salicylate inhibition phenotype to wildtype SlyA, indicating that allosteric inhibition does not occur by disulfide bond formation between cysteine residues *in vivo*. We subsequently generated a chromosomal C81S mutant to determine whether SlyA is directly inhibited by Cu^2+^. Transcription of *pagC* in strains encoding wildtype or C81S *slyA* was reduced approximately two-fold after the addition of 100µM CuCl_2_ to the growth medium (Figure 4B). Although this difference was statistically significant (p<0.001), it was also observed in the mutant strain encoding C81S SlyA, indicating that this modest effect is not due to disulfide-dependent tetramerization as proposed for MarR (15). Furthermore, modulation by copper does not appear to be biologically significant in comparison to the 350-fold effect of salicylate. Cu^2+^-mediated derepression of the *marRAB* operon was confirmed under comparable conditions by measuring *marA* expression, which increased approximately 7.5-fold (Figure 4C).

### Evolutionary analysis of the SlyA TF lineage

To identify other variables that may have contributed to the evolution of the SlyA lineage, we performed an evolutionary analysis of species representing 60 genera of *Enterobacteriaceae*, including *candidatus* organisms for which genomic data was available. SlyA is strongly conserved, with orthologs found in 55 organisms, suggesting that SlyA plays a central role in enterobacterial regulatory circuitry. Genera lacking identifiable SlyA orthologs include *Buchnera, Hamiltonella, Samsonia, Thorsellia*, and *Plesiomonas*, which are only distantly related to *Salmonella* and *Escherichia* in comparison to other members of the *Enterobacteriaceae*. Notably, we were also unable to identify an *hns* ortholog in *Plesiomonas.* Phylogenetic analysis reveals five clusters of SlyA orthologs (Figure 5), with the general structure of the phylogram resembling the genomic tree for *Enterobacteriaceae* (37). SlyA orthologs in enteric pathogens including *E. coli, S.* Typhimurium, and *S. flexneri* form cluster I, while cluster II is composed of a more heterogeneous group of organisms, including plant pathogens such as *D. dadantii*, insect endosymbionts like *S. glossinidius*, and more distantly related pathogens such as *Y. pseudotuberculosis*. SlyA orthologs in clusters I and II are known to function as pleiotropic regulators, despite significant divergence from the other clusters (28, 31). Clusters III and IV are comprised of environmental organisms such as the plant-associated species *Pantoea agglomerans* and *Phaseolibacter flectens*, while cluster V contains hydrogen-sulfide producing bacteria such as *Pragia fontium*. This degree of conservation suggests that SlyA diverged from the greater MarR family of TFs prior to the divergence of *Enterobacteriaceae* from other bacteria. Although not strongly conserved outside of *Enterobacteriaceae*, SlyA orthologs can be detected in selected species throughout the Gammaproteobacteria, indicating that the lineage is ancient. However, the *slyA* gene typically exhibits an AT-content (51% in *S*. Typhimurium) marginally higher than that of the chromosomal average (48% in *S*. Typhimurium), suggesting that it may have been duplicated through an ancient transfer event.

**Figure 5.**
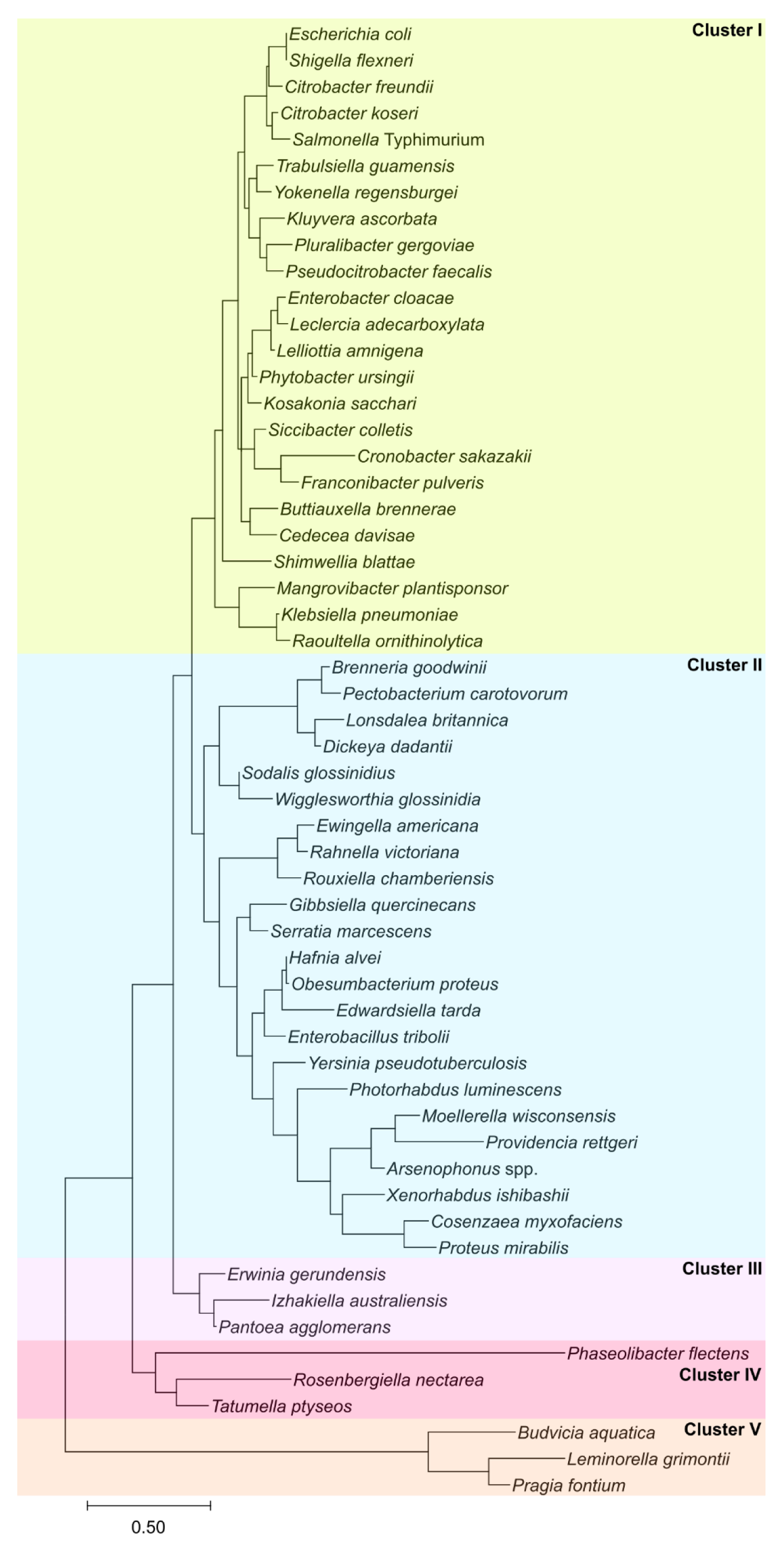
The SlyA TF lineage in *Enterobacteriaceae*. The evolutionary history of the SlyA TF lineage was inferred using the Maximum Likelihood method and the Le-Gascuel model (85) with Mega X software (86). The tree is drawn to scale with the branch length representing the number of substitutions per site. The *slyA* genes have evolved in five major phylogenetic clusters: cluster I comprises most enteric species, including *S*. Typhimurium and *E. coli*; cluster II is heterogeneous, containing more distantly related pathogenic species from *Proteus* and *Yersinia*, as well as endosymbionts like *Sodalis*; clusters III and IV contain several plant-associated bacteria, such as *Pantoea agglomerans* and *Phaseolibacter flectens*; cluster V is comprised of hydrogen sulfide-producing bacteria. (A multiple sequence alignment of selected SlyA orthologs can be found in Figure 5 - figure supplement 1).

To determine if allosteric inhibition is likely to be conserved throughout the SlyA lineage, we aligned the sequences of SlyA orthologs from different bacterial species to analyze the conservation of residues involved in salicylate binding (Table S3). Cluster I exhibited nearly complete conservation of the salicylate-binding residues, with only four of 24 genomes encoding polymorphisms. Cluster II exhibited the greatest variation, with 16 of 23 genomes encoding polymorphisms, including 10 which encoded the polymorphism H38Y. We identified only one polymorphism in site I in six genomes, a T32I substitution, suggesting that site I is particularly important for SlyA function. As further evidence of the importance of site I, a T66A mutation abrogated allosteric inhibition of SlyA. The central residues of site II (R14, W16, R17, and W34) are similarly conserved suggesting that these residues are also important for SlyA function. We were able to mutate two of these residues (Figure 4A), R14 and W34, and found that W34 is also involved in allosteric inhibition. Individual polymorphisms such as T32I and H38Y appear to have evolved independently in multiple clusters, suggesting purifying selection, although their effect on SlyA activity is currently unknown. Alignment of SlyA protein sequences from representative species of each of the five clusters revealed that most sequence variation occurs in the carboxyl-terminus oligomerization domain (Figure 5 - figure supplement 1), suggesting that ligand sensitivity and DNA binding are conserved features of the SlyA lineage.

As the ancestral function of MarR TFs is the negative regulation of genes encoding drug efflux pumps, a function that is conserved in *S*. Typhimurium, we examined *slyA* orthologs throughout the *Enterobacteriaceae* for linkage to flanking genes of known or hypothetical function: *slyB*, which encodes an putative outer membrane lipoprotein with no observed phenotype, *ydhJ* which encodes a hemolysin D homolog, and *ydhK*, encoding the efflux pump (Figure 6A, B). It should be noted that *ydhJ* also exhibits homology to *emrA*, which encodes an antimicrobial efflux pump-associated protein linked to and regulated by another MarR family TF called MprA or EmrR (38, 39). We found that cluster I exhibited the strongest linkage, with 100%, 95%, and 100% linkage to *ydhJ, ydhK*, and *slyB* respectively, while cluster V, containing the hydrogen sulfide-producing bacteria, exhibited no linkage to *slyA* for any of the genes examined. Outside of cluster V, *slyB* was strongly associated with *slyA*, exhibiting 76% linkage overall. However, the *ydh* operon was not strongly linked outside of cluster I, exhibiting 26%, 33% and 33% linkage in clusters II, III, and IV respectively. In contrast, although *marR* was absent from 26 of the enterobacterial species examined, *marR* was linked to *marAB* in every *marR-*carrying species. This suggests that the primary function of MarR is to regulate the *marRAB* operon, and MarR consequently does not play an important role in enterobacterial regulatory circuitry. In contrast, the loss of the ancestral linkage between *slyA* and *ydhJ* outside of cluster I suggests that the SlyA lineage has evolved to serve a distinct function.

**Figure 6.**
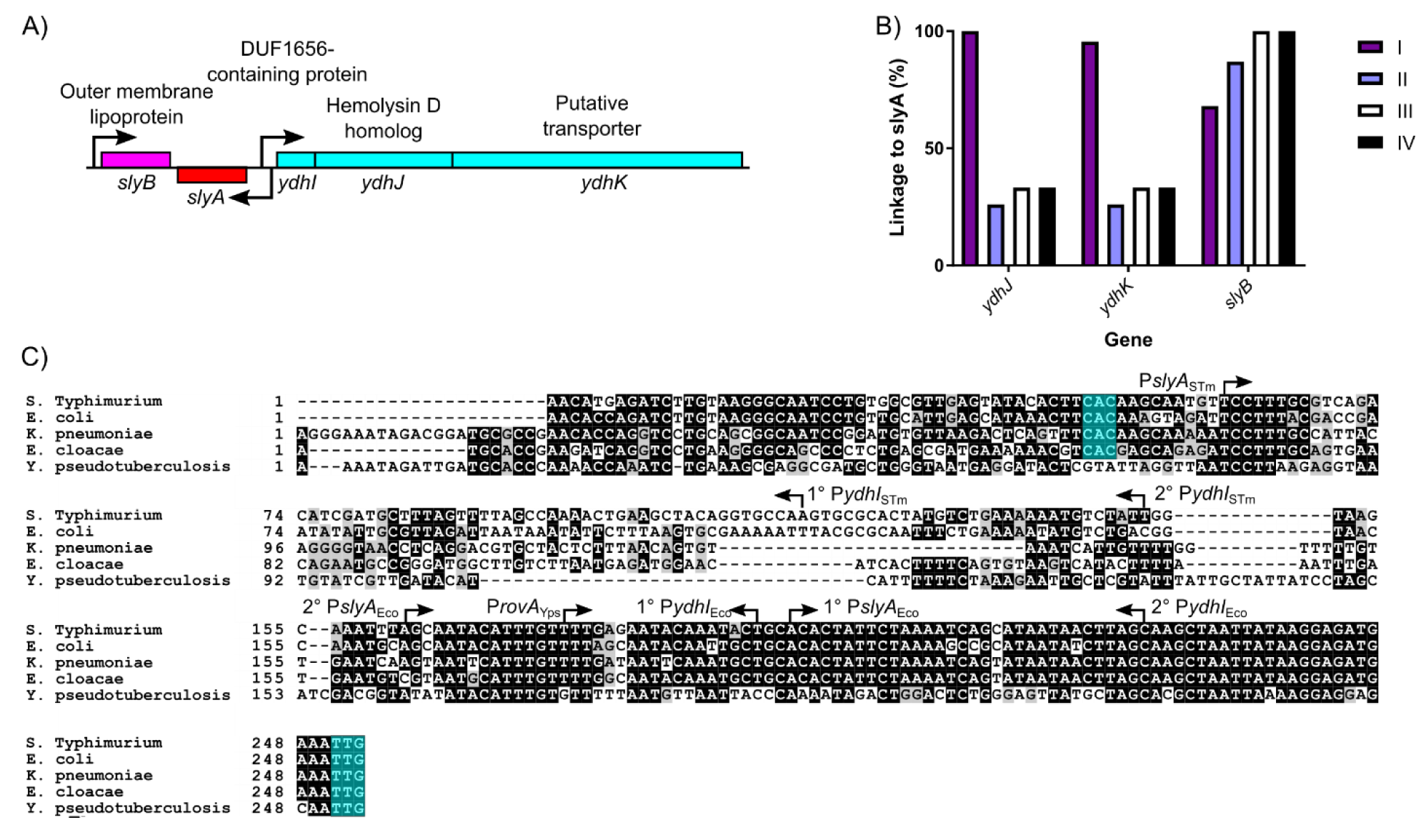
Genetic linkage and *cis*-regulatory evolution of *slyA* in *Enterobacteriaceae*. The *slyA* region of *S.* Typhimurium and closely related enteric species is diagrammed in (A). Arrows indicate TSSs, and boxes represent protein coding sequences. The *ydhIJK* efflux pump operon is transcribed divergently from *slyA*. A protein of unknown function is encoded by *ydhI,* and *ydhJ* and *ydhK* encode a hemolysin D/EmrA homolog and a transporter protein, respectively. *slyB* encodes a putative outer membrane lipoprotein, transcribed convergently to *slyA*. The linkage of *slyA* to *slyB, ydhJ*, and *ydhK*, is determined for all of the species from clusters I-IV shown in Fig. 5 (B). Species from cluster V do not exhibit linkage to these genes and are not shown. A multiple sequence alignment of the *slyA* promoter region from the closely-related species *S*. Typhimurium, *E. coli, K. pneumoniae*, and *E. cloacae*, as well as more distantly-related *Y. pseudotuberculosis* (C). Arrows indicate the position and orientation of previously characterized TSSs (30, 55, 57). Where multiple TSSs have been identified, the primary (1°) and secondary (2°) start sites are indicated. The *slyA* (positions 248-250) and *ydhI* (positions 47-70, dependent on the species) start codons are highlighted in teal when present. (Alignments of this region from representative species of *Salmonella, Escherichia*, and *Yersinia* are shown in Figure 6 - figure supplement 1. A phylogenetic analysis of the *slyA* promoter region of a subset of genera is shown in Figure 6 - figure supplement 2.)

The *ydhIJK* operon does not appear to have been exchanged for another drug efflux system. (Figure 6C). Sequences homologous to the proximal portion of the *ydhI* gene are detectable even in species that have not retained functional coding sequences (*e.g., Y. pseudotuberculosis*). However, a 100 bp segment of the *ydhI-slyA* intergenic region, beginning approximately 95 bp upstream of the *slyA* start codon, has undergone extensive mutation (Figure 6C), suggesting *in cis* evolutionary adaptation in the *slyA* lineage of TFs. We examined the regions upstream of *slyA* in several related species of three genera, *Escherichia, Salmonella*, and *Yersinia*, to better understand the recent evolution of this region, and observed that the *ydhI-slyA* intergenic regions exhibit considerable divergence between genera and conservation within genera (Figure 6 - figure supplement 1). A comparison between the *ydhI-slyA* intergenic regions of *Escherichia coli* and *Salmonella* Typhimurium with those of *Klebsiella pneumoniae* and *Enterobacter cloacae* revealed similar degrees of divergence between each genus, suggesting that an ancestral allele can no longer be defined. A phylogenetic analysis of *slyA* upstream regions in a representative subset of the species described above demonstrated extensive variation throughout the *Enterobacteriaceae* (Figure 6 - figure supplement 2). However, this variation did not always correlate with SlyA coding region clusters. Although the intergenic regions of cluster I are closely related and may have co-evolved with their respective coding sequences, the other clusters are more variable, and some intergenic regions may have evolved independently of their coding sequences.

Examination of the *ydhI-rovA* region in *Yersinia spp.* reveals that *rovA* in *Y. pseudotuberculosis* (Figure 6C) and *Y. pestis* (Figure 6 - figure supplement 1) is not linked to a functional *ydhIJK* operon, whereas *ydhIJK* is retained in *Y. enterocolitica*. This indicates that *ydhIJK* was not lost by the former species until after the divergence of *Yersinia* from other *Enterobacteriaceae* and suggests that other species of *Enterobacteriaceae* are also likely to have lost *ydhIJK* recently, as each became adapted to its own specific niche. This represents an example of parallel evolution, with multiple species independently losing *ydhIJK* to accommodate *in cis* evolution as their respective SlyA orthologs adapted to new roles.

### Functional characterization of a distantly related SlyA ortholog

To understand how the SlyA lineage adapted to its emergent role as an important pleiotropic regulatory protein in the *Enterobacteriaceae*, we compared the functions of SlyA from *S*. Typhimurium (SlyA_STM_) and RovA of *Y. pseudotuberculosis* and *Y. enterocolitica,* a relatively divergent ortholog that also functions as a counter-silencer (28-30, 40). RovA of *Y. pseudotuberculosis* exhibits 76% identity with SlyA_STM_ (Figure 5 - figure supplement 1). RovA is essential for virulence in *Y. enterocolitica*, and *Y. pestis* (28, 41) and has been suggested to function both as an activator, interacting with RNAP (42), and as a counter-silencer, alleviating H-NS-mediated repression (29, 30, 40). Notably, direct activation has not been demonstrated for any other member of the SlyA lineage, and the most direct evidence to suggest that RovA functions as an activator is derived from IVT studies using small linear fragments of DNA as a template (42). We have previously shown that small linear fragments are not necessarily representative of physiological regulatory events in intact cells and can generate spurious results in IVT assays (17). Genetic analysis of the *inv* gene, which is positively regulated by RovA, in an *hns* mutant strain of *E. coli* also suggested that RovA functions as both a counter-silencer and an activator. However, these experiments failed to consider the potential contribution of the H-NS paralog, StpA, which is up-regulated in *hns* mutants and can provide partial complementation (43, 44), which complicates the interpretation of regulatory studies in an *hns* mutant strain.

We attempted to corroborate the published findings by performing IVT analysis of *inv* expression in the presence of RovA. However, expression of *inv* exhibited only a very modest (∼1.5-fold) increase following the addition of RovA (Figure 7A), which became saturated at a 20 nM concentration; these results are not supportive of direct *inv* activation by RovA. In contrast, RovA was confirmed to function as an auto-repressor, like other MarR family TFs, as IVT analysis demonstrated a 4-fold decrease in *rovA* expression following the addition of RovA protein, with the effect reaching saturation at a 500 nM concentration (Figure 7B). RovA has also retained the ability to respond to salicylate (Figure 7C), despite the presence of an H38Y substitution in the second salicylate-binding pocket (Table S3), as 5mM salicylate completely inhibited RovA-mediated repression, similar to our observations with SlyA. It is also notable that RovA has retained salicylate sensitivity despite the loss of the linked YdhIJK efflux pump in *Y. pseudotuberculosis*. To prove that RovA functions as a counter-silencer in *Y. pseudotuberculosis*, we measured *inv* expression in wild-type and *rovA* mutant strains expressing H-NST from enteropathogenic *E. coli* (H-NST_EPEC_). Mutations in *hns* cannot be generated in *Yersinia* as *Yersinia* spp. carry a single essential *hns* gene, unlike many other members of the *Enterobacteriaceae* which encode *hns*-like genes such as *stpA*, which are able to partially compensate for the loss of *hns*. H-NST_EPEC_ is a truncated *hns* homolog that has been demonstrated to function as a dominant negative form of H-NS by binding and inhibiting the activity of wild-type H-NS protein (45). The expression of *inv* in *rovA* mutant bacteria was approximately four-fold lower than in wild-type cells (Figure 7D). However, *inv* expression was fully restored upon inhibition of H-NS by *hnsT*_EPEC_, demonstrating that RovA functions solely as a counter-silencer of the *inv* gene. The *rovA* and *slyA* genes are capable of complementing each other in both *S.* Typhimurium and *Y. pseudotuberculosis* when expressed *in trans* (Figure 7E, F), up-regulating both *inv* and *pagC*, further indicating that RovA functions as a counter-silencer, like SlyA. Together, these observations demonstrate that RovA has retained the ancestral characteristic of environmentally-responsive repression exhibited by other members of the MarR TF family, despite being one of the most divergent members of the SlyA lineage. However, in contrast to MarR TFs outside of the SlyA lineage, it is also able to function as a counter-silencer of horizontally-acquired genes, as exemplified by *inv*.

**Figure 7.**
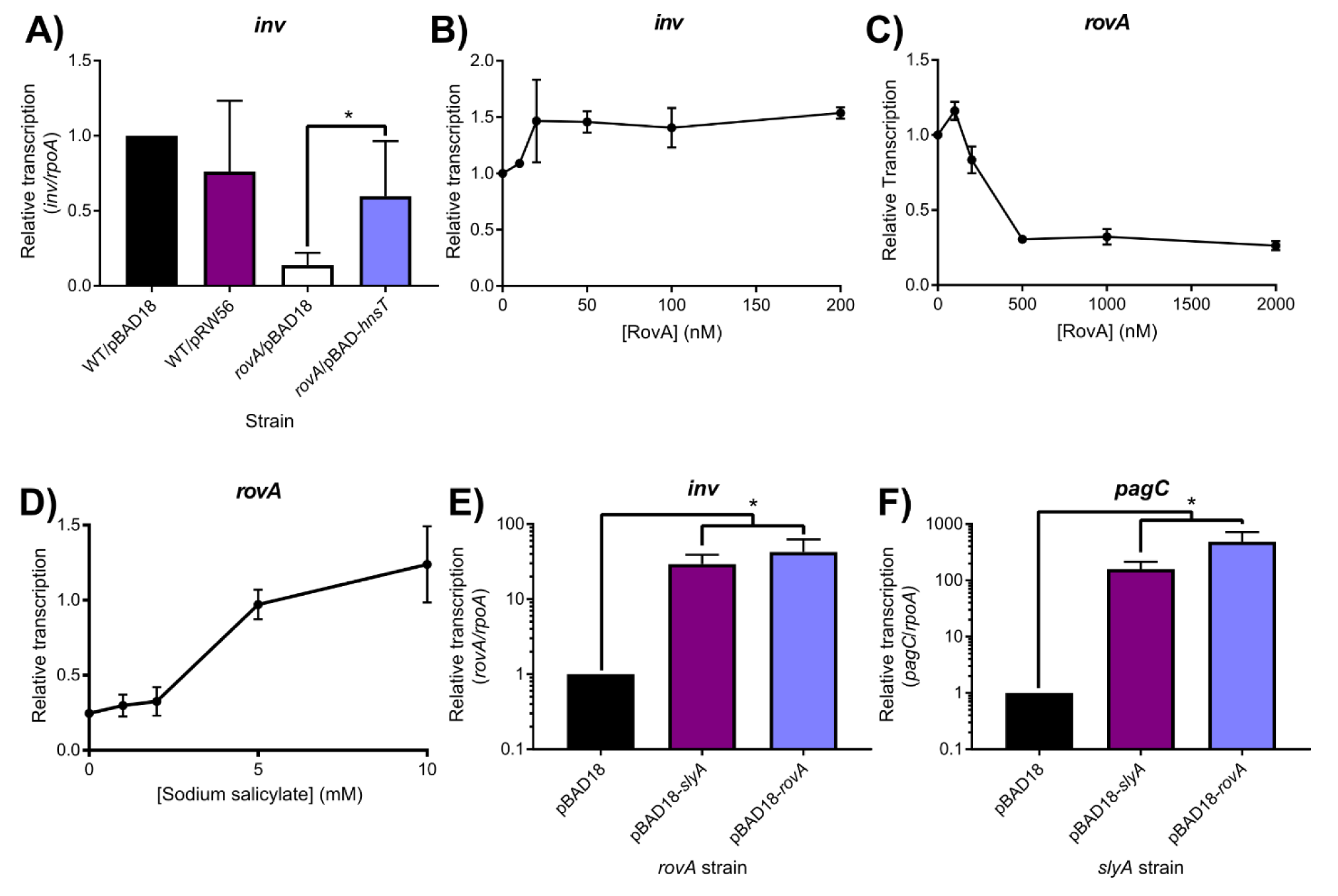
Environmentally-responsive repressing and counter-silencing functions are conserved in the SlyA TF lineage. To determine whether RovA functions as a counter-silencer in *Y. pseudotuberculosis*, transcription of the Rov-regulated *inv* gene was quantified by qRT-PCR in wildtype or *rovA* mutant strains expressing *hnsT*_*EPEC*_, which inhibits H-NS (A). Although *inv* expression is decreased in a *rovA* mutant strain, expression is fully restored by the inhibition of H-NS, indicating that RovA functions as a counter-silencer. IVT assays of the *inv* regulatory region in the presence of increasing RovA concentrations detected only a minimal impact on transcription levels, suggesting that RovA is unlikely to function as a classical activator (B). IVT assays of *rovA* in the presence of increasing RovA concentrations demonstrate that RovA functions as an autorepressor (C). RovA remains sensitive to inhibition by small aromatic carboxylate molecules, as increasing concentrations of sodium salicylate inhibited RovA-mediated *rovA* repression in IVT assays. Reciprocal complementation studies were performed by providing either *rovA* or *slyA in trans* expressed from the arabinose-inducible pBAD18 vector in *rovA* or *slyA* mutant *Y. pseudotuberculosis* and *S*. Typhimurium strains, respectively, and measuring transcription of *inv* and *pagC* via qRT-PCR. Transcript levels are normalized to *rpoA* in both species, and data represent the mean ± SD; n=3. Asterisks indicate P < 0.05.

### SlyA counter-silencing requires high expression levels

It is interesting to note that SlyA does not play a major regulatory role in *Escherichia coli,* the most well-studied member of the *Enterobacteriaceae*. SlyA is only known to regulate two *E. coli* genes, *hlyE* and *fimB* (46-48), despite exhibiting a high degree of homology (91% identity) to SlyA in *S.* Typhimurium. To better understand the different roles of SlyA in the regulatory hierarchy of *E. coli* and *S.* Typhimurium, we performed a comparative genetic analysis of the *S*. Typhimurium (*slyA*_STm_) and *E. coli* (*slyA*_Eco_) alleles. In allelic exchange experiments, the *slyA*_STm_ coding sequence was swapped with *slyA*_Eco_ to determine the effect on the expression of the counter-silenced *S.* Typhimurium *pagC* gene (Figure 8A). We observed that SlyA_Eco_ is able to counter-silence the expression of *pagC* similarly to SlyA_STm_, suggesting that the diminished role of SlyA in *E. coli* is not attributable to differences in protein sequence. This suggested that the importance of different SlyA lineage proteins within their respective regulatory networks may be the result of the different *in cis* evolutionary pathways identified in our phylogenetic analysis (Figure 6 and Figure 6 - figure supplement 1), resulting in differences in levels of expression. To test this hypothesis, *slyA* intergenic region-ORF chimeras were constructed and assessed for their ability to counter-silence *pagC*. To avoid potentially confounding results due to multiple transcriptional start sites (TSSs), we exchanged the intergenic regions beginning immediately upstream of each start codon. Counter-silencing was assessed in a *slyA* mutant carrying pKM05, a plasmid with the *slyA*_STM_ ORF transcribed by the *E. coli* promoter (P*slyA*_Eco_), or pKM07, a plasmid with the *slyA*_STM_ ORF transcribed by its native promoter (P*slyA*_STM_). Although *pagC* expression was similar with either construct under non-inducing conditions, expression was approximately 10-fold lower with *slyA* expressed from P*slyA*_Eco_ under inducing conditions (Figure 8B), suggesting that the diminished role of SlyA in *E. coli* is at least partially attributable to differences in *slyA* expression levels in *S*. Typhimurium and *E. coli*. Although *slyA* expression driven by P*slyA*_Eco_ is only slightly lower under non-inducing conditions, it is approximately 3-fold greater when driven by P*slyA*_STM_ under inducing conditions (Figure 8C), suggesting that P*slyA*_Eco_ is less responsive to environmental signals associated with virulence gene expression in *Salmonella*, and thus is unable to regulate *slyA* expression in a manner appropriate for counter-silencing. Parallel experiments in *Y. pseudotuberculosis* revealed that *rovA* is also more strongly expressed from its native promoter (P*rovA*_Yps_) than from a chimera expressing *rovA* from P*slyA*_Eco_ (Figure 8 - figure supplement 1), despite the fact that P*rovA*_Yps_ has diverged much more significantly from both P*slyA*_Eco_ and P*slyA*_STm_ than P*slyA*_Eco_ and P*slyA*_STm_ have diverged from each other (Figure 6 - figure supplement 1).

**Figure 8.**
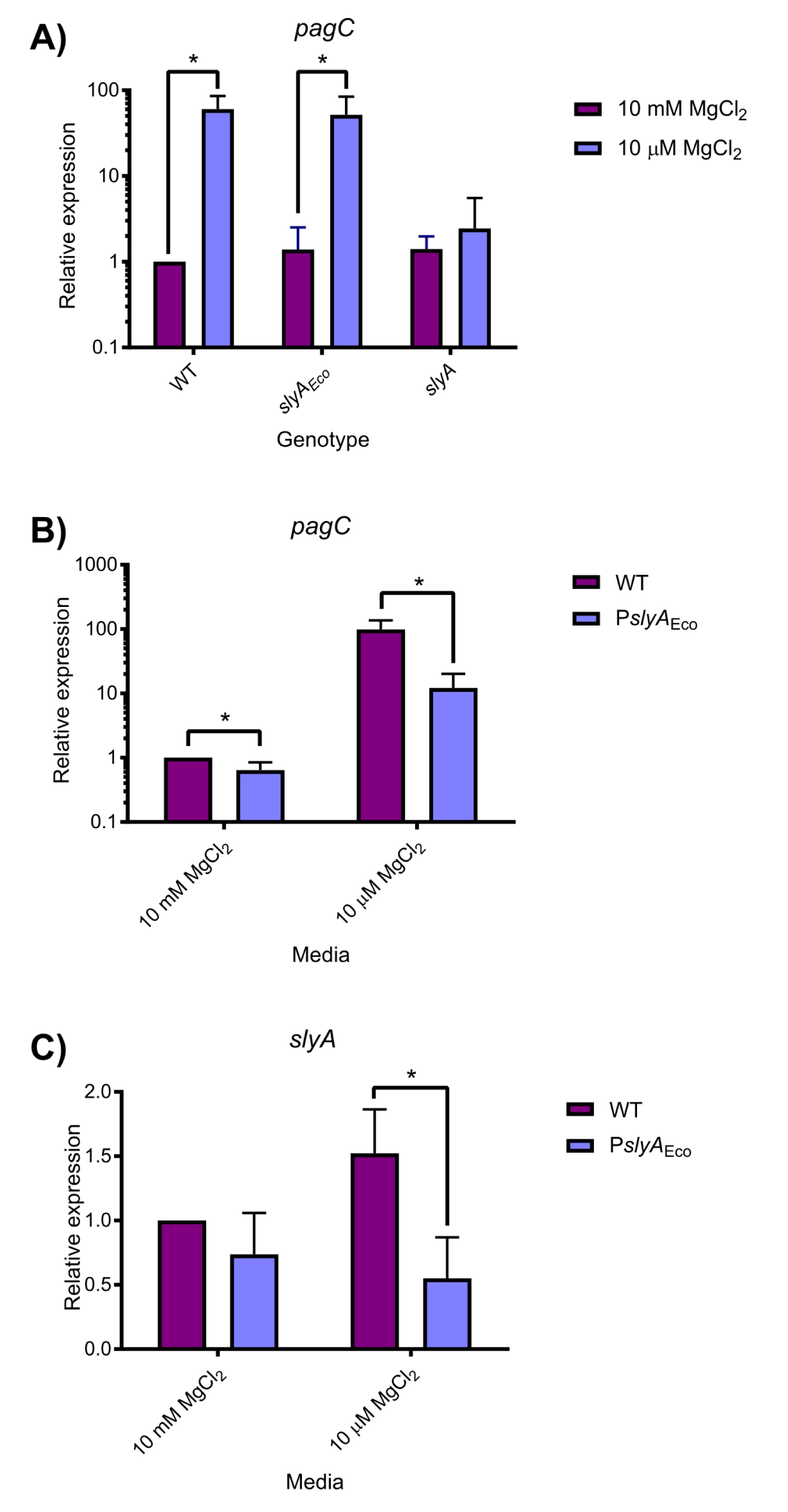
*S*. Typhimurium SlyA is a more effective counter-silencer than *E. coli* SlyA due to higher expression levels. Expression of the SlyA counter-silenced *pagC* gene was measured by qRT-PCR in *S.* Typhimurium strains encoding wildtype *slyA, slyA*_*Eco*_, or mutant *slyA* under either inducing (10 µM MgCl_2_) or non-inducing (10mM MgCl_2_) conditions (A). Expression of *pagC* (B) and *slyA* (C) was compared in *S*. Typhimurium strains in which *slyA* transcription was driven by either the wildtype (WT) or *E. coli* (P*slyA*_*Eco*_) *slyA* promoter. Transcript levels are normalized to *rpoD* and data represent the mean ± SD; n=3. Asterisks indicate P ≤ 0.05. (A comparative analysis of a P*slyA*_Eco_-*rovA* chimera is shown in Figure 8 - figure supplement 1.)

## Discussion

The ancient MarR family of transcription factors is represented throughout the bacterial kingdom and in many archaeal species (36). This study sought to understand how the SlyA/RovA lineage of MarR TFs in the *Enterobacteriaceae* evolved to acquire the novel function of counter-silencing. Our observations demonstrate that SlyA proteins have retained vestiges of their ancestral functions, e.g., the environmentally-responsive repression of small molecule efflux systems, while acquiring an ability to act as pleiotropic counter-silencers of horizontally-acquired genes. The latter role has been facilitated by the parallel evolution of cis-regulatory elements that support higher levels of gene expression.

SlyA, like other MarR family TFs, is subject to allosteric inhibition by small aromatic carboxylate compounds such as salicylate, which bind and stabilize the SlyA dimer in a conformation unfavorable for DNA binding (Figure 2 and Figure 2 - figure supplement 1). However, the specific structure and arrangement of the effector binding sites varies significantly among the MarR family. Our structural data indicate that SlyA binds a total of six salicylate molecules per dimer, although two of these are not likely to be biologically relevant (Figure 3). Although the general structure and architecture of the salicylate-SlyA complex is similar to complexes formed by other MarR TFs, including MarR (8) and MTH313 from *Methanobacterium thermautrophicum* (49), the various TFs differ significantly in their specific interactions with salicylate. MarR binds a total of four salicylate molecules per dimer, whereas MTH313 binds only two (Figure 3D, E). All four salicylate-binding sites in MarR flank the wHTH domain and are partially exposed to solvent, whereas none of these sites corresponds to the salicylate binding sites of SlyA, potentially accounting for the significant differences in affinity for aromatic carboxylates between the two proteins. MTH313 binds salicylate asymmetrically at sites similar to sites I and II of SlyA, binding only one salicylate molecule per monomer. It is possible that the structural variability in the MarR family with respect to salicylate binding reflects an inherent evolutionary flexibility and ligand promiscuity. The MarR family sensor region may have evolved to interact with a variety of small molecules, and salicylate may simply represent a promiscuous probe for potential interactions. Recent studies have suggested that the true ligand of MarR may be copper, liberated from membrane-associated proteins during oxidative stress induced by xenobiotic agents. Copper reportedly oxidizes a cysteine residue (C80) to promote disulfide bond formation between MarR dimers, resulting in subsequent tetramerization and inhibition of DNA binding (15). As almost all characterized MarR TFs have a reactive cysteine residue corresponding to a region near Site II of SlyA, we considered that cysteine oxidation, which has also been described in the MarR family TF OhrR (50), might occur in the SlyA lineage as well. However, mutation of C81, the lone cysteine residue in SlyA_STm_, had a negligible effect on SlyA activity in both the presence and absence of salicylate. The addition of copper to the growth medium had a minimal impact on SlyA activity and was unaffected by mutation of C81, indicating that cysteine oxidation is not a general mechanism for the allosteric inhibition of MarR family TFs. Furthermore, the high affinity of SlyA for aromatic carboxylates (Figure 2 - figure supplement 2) and the conservation of binding pocket residues (Table S3) reported in this study suggest that environmental sensitivity and allosteric inhibition are a conserved feature of SlyA activity in their new regulatory role.

A phylogenetic analysis of SlyA orthologs in *Enterobacteriaceae* demonstrates that the SlyA lineage is strongly conserved, even in endosymbiotic species exhibiting significant genome loss (Figure 5 and Figure 5 - figure supplement 1). This suggests that SlyA has a central and essential role in the transcriptional regulatory networks of these species. For example, a recent analysis of the endosymbiont *W. glossinidia* found that *slyA* is subject to significant evolutionary constraints (51). This conservation does not appear to be due to its role as a regulator of antimicrobial resistance, as *slyA* does not exhibit significant linkage to the efflux pump operon *ydhIJK*, outside of the enteric pathogens in cluster I (Figure 6). Rather, SlyA proteins appear to function predominantly as pleiotropic counter-silencers, facilitating the integration of horizontally-acquired genes, including virulence genes, into existing regulatory networks. A general counter-silencing role has been suggested in multiple enterobacterial species including *E. coli* (19, 48), *Salmonella* (17, 20, 21) and *Shigella* (32) in cluster I, and *Yersinia* (31, 52), *Serratia*, and *Pectobacterium* (*Erwinia*) (25) in cluster II. Although similar evidence does not currently exist for the endosymbionts of cluster II, this may simply reflect the limited genetic analyses that have been performed in these species.

To understand the evolutionary adaptations to accommodate counter-silencing by the SlyA/RovA lineage, we compared three orthologs, *slyA*_STM_, *slyA*_Eco_ and *rovA*. The *slyA*_STM_ and *rovA* genes are essential for *Salmonella* and *Yersinia* virulence, whereas *slyA*_Eco_ plays a negligible role in *E. coli*, despite exhibiting a much higher degree of similarity to *slyA*_STM_ than does *rovA*. These differing roles are attributable to differences in expression, as SlyA_Eco_ is able to function as a counter-silencer in *S*. Typhimurium when its expression is driven by the *S*. Typhimurium promoter (Fig. 6). We also observed that *slyA*_Eco_ transcription is significantly reduced in medium with low Mg^2+^ concentrations, a condition associated with the *Salmonella*-containing vacuole (SCV) of macrophages, in which SlyA cooperates with the response regulator PhoP to counter-silence virulence genes necessary for intracellular survival (17). This suggests that the diminished role of *slyA*_Eco_ in *E. coli* may result from its inability to respond to appropriate environmental cues. This is further reinforced by the observation that RovA of *Yersinia* spp. is able to function as a counter-silencer in *S*. Typhimurium when expressed *in trans* from an inducible promoter, and that the *rovA* promoter of *Y. pseudotuberculosis* drives *rovA* transcription more strongly than P*slyA*_Eco_ in a chimeric strain (Fig. S7). This suggests that evolution of the SlyA-RovA lineage *in trans*, particularly DNA binding specificity, has played only a minor role since the divergence of the *Enterobacteriaceae*. This is further supported by as comparison of distantly-related SlyA orthologs, which exhibit most sequence divergence in the C-terminal oligomerization domain and not the N-terminal region containing the wHTH (Figure 5 - figure supplement 1). We conclude that the ability of a given SlyA ortholog to serve as a counter-silencer is contingent on its level and pattern of expression, which may be the product of both transcriptional and post-transcriptional activity, as mutations altering TSS position will subsequently alter the 5’ untranslated region of the *slyA* transcript. Notably, another group recently demonstrated a significant expansion of the SlyA regulon in *E. coli* when *slyA*_Eco_ is overexpressed, with 30 operons exhibiting regulation by SlyA, 24 of which are also repressed by H-NS (34), indicating that SlyA_Eco_ is capable of functioning as a counter-silencer but is not expressed under the appropriate conditions. In *S*. Typhimurium, the appropriate conditions are those associated with the SCV. However, SlyA orthologs in other species such as the plant pathogens and endosymbionts of cluster II are likely to require expression under vastly different conditions corresponding to their environmental niches. The loss of the divergently-transcribed *ydhIJK* operon in *Y. pseudotuberculosis* and other enteric species may be a consequence of genetic alterations to enhance *rovA* expression as well as the redundancy of drug efflux pumps. An additional possibility is that enhanced *slyA/rovA* expression might result in hyper-repression of *ydhIJK*, which would negate the usefulness of the pump to the cell. Even the plant pathogens such as *D. dadantii* and *P. carotovorum*, which are most likely to encounter small phenolic compounds in the plant environment (53), have failed to retain *ydhIJK*. It is also notable that the *slyA* orthologs in the four species (*E. coli, S.* Typhimurium, *Y. enterocolitica, Y. pseudotuberculosis*) in which the transcriptional start sites have been characterized initiate transcription at different positions (30, 54-57), indicating that each species has evolved its *cis*-regulatory circuit independently (Figure 6).

Previous studies have demonstrated that regulatory evolution can promote adaptation to new niches (58). The SlyA/RovA TF lineage provides a unique example of parallel regulatory evolution to achieve a common functional objective. Throughout the *Enterobacteriaceae*, the associated *ydhIJK* pump genes have been repeatedly lost, yet their regulators have been retained, presumably to facilitate the evolution of an appropriately responsive regulatory circuit to enable counter-silencing. This suggests that SlyA/RovA proteins possess intrinsic features that predispose them for a counter-silencing role, perhaps the abilities to respond to environmental stimuli and to recognize a variety of AT-rich target DNA sequences. Studies are underway to characterize these features and to determine their contribution to the evolutionary capacity of *Enterobacteriaceae*.

## Materials and methods

### Bacterial strains and general reagents

All oligonucleotides and plasmids used in this study are described in Table S4 and Table S5, respectively. Unless otherwise indicated, bacteria were grown in Luria Bertani (LB) broth with agitation. *Salmonella enterica* serovar Typhimurium strains were constructed in the ATCC 14028s background and grown at 37°C, unless otherwise indicated. The 14028s *slyA* mutant strain was described previously (17). *Yersinia pseudotuberculosis* YPIII (59) and YP107 (60) (a gift from P. Dersch, Helmholtz Centre for Infection Research) were used as the wildtype and *rovA* strains, respectively, and grown at 24°C. A *ydhIJK* deletion mutant was constructed using the λ-Red recombinase system (61) and oligonucleotides WNp318 and WNp319. A *slyA ydhIJK* strain was generated by introducing the *slyA::Cm* cassette from 14028s *slyA::Cm* (17) to 14028s *ydhIJK* via P22HTint-mediated transduction. To exchange the wild-type *S*. Typhimurium *slyA* coding sequence with the *E. coli* allele, *S.* Typhimurium *slyA* was replaced with a *thyA* cassette, via FRUIT (62), using the oligonucleotides STM-slyA-targ-F and STM-slyA-targ-R. The *slyA* coding sequence from *E. coli* K-12 was then amplified using the primers STM-Eco-slyA-F and STM-Eco-slyA-R, which include 40 and 41 bases, respectively, from the regions flanking the *S*. Typhimurium *slyA* coding sequence. This fragment was electroporated into the *slyA::thyA* mutant strain and the resulting transformants were plated on minimal media containing trimethoprim, as described for the FRUIT method (62), to select for replacement with the *E. coli slyA* allele.

The chromosomal C81S *slyA* mutation (TGC→AGC) was generated by cloning a 1 kb fragment encoding the C81S mutation into the suicide vector pRDH10 (63) using Gibson Assembly (New England Biolabs, Ipswich MA). The Gibson assembly reaction included PCR products generated with primers JKP736/JKP737 and JKP738/JKP739 and genomic *S*. Typhimurium DNA as well as *Bam*HI-digested pRDH10 to generate pJK723. For integration of *slyA* C81S into the chromosome, S17-1Δλpir (64)/pJK723 was mated with *S.* Typhimurium 14028s/pSW172 (65) and plated onto LB+20 µg ml^-1^ chloramphenicol and incubated at 30°C overnight. Chloramphenicol- and carbenicillin-resistant (Cm^r^ Carb^r^) colonies that were isolated represented a single crossover event of pJK723 (Cm^r^) plasmid into the *slyA* region of *S.* Typhimurium 14028s (Carb^r^). Selection for the second crossover event to replace wild-type *slyA* with a *slyA* C81S mutation was performed by plating 0.1ml of an overnight culture of Cm^r^ Carb^r^ colony onto LB+5% sucrose plates and incubating at 30°C overnight. Colonies were then streaked onto LB plates at 37°C and putative 14028s *slyA* C81S colonies confirmed by DNA sequencing.

### Cloning

pSL2143 was generated by amplifying the *slyA*_STm_ region, including its native promoter, with primers slyAcomp-F and slyAcomp-R, and ligating into pWSK29 (66) digested with *Eco*RV. The G6A, S7A, A10P, R14A, W34A, H38A, T66A and C81A mutants were generated with their respective mutagenic primer pairs (Table S4) using the QuikChange XL Site-Directed Mutagenesis Kit (Agilent Technologies, Santa Clara CA) according to the manufacturer’s protocol. A 2249 bp region containing both *slyA* and *ydhI* was amplified by PCR using the primers SlyAreg-F and SlyAreg-R. The resulting fragment was digested with *Bam*HI and *Hin*dIII and ligated into the low-copy IVT scaffold vector pRW20 (17) to generate pRW39. An IVT target containing the 2379 bp *rovA* region from *Y. pseudotuberculosis* was generated using the primers BamHI-rovA-F and EcoRI-rovA-R. The resulting fragment and pRW20 were both digested with *Bam*HI and *Eco*RI and ligated together to construct pRW54. An IVT target containing the 2708 bp *inv* region from *Y. pseudotuberculosis* was generated via PCR using the primers BamHI-inv-F and EcoRI-inv-R. The resulting fragment and pRW20 were both digested with *Bam*HI and *Eco*RI and ligated together to construct pRW55. To perform *rovA* and *slyA* complementation studies, both genes were cloned into the arabinose-inducible expression vector pBAD18 (67). The *rovA* gene was amplified using the oligonucleotides EcoRI-rovA-F and KpnI-rovA-R. The *slyA* gene was amplified using the oligonucleotides EcoRI-slyA-F and KpnI-slyA-R. Both fragments were digested with *Eco*RI and *Kpn*I, and ligated into pBAD18, generating pRW58 (*slyA*) and pRW59 (*rovA*). pRW60 was constructed by cloning an *N*-terminal 6×His-tagged copy of *rovA* into pTRC99a (68). The *rovA* gene was amplified from YPIII genomic DNA by PCR using the oligonucleotides 6HisRovA-F and 6HisRovA-R. The resulting fragment and pTRC99a were digested with *Nco*I and *Bam*HI, and ligated together. An *E. coli* P_*slyA*_-S. Typhimurium *slyA* coding sequence chimera was constructed using overlapping PCR. The *slyA-ydhIJK* intergenic region was amplified from *E. coli* K12 genomic DNA using primers KMp178 and KMp206, and from *S*. Typhimurium 14028s using oligonucleotides KMp177 and KMp206. The corresponding *slyA* ORF from *S*. Typhimurium was amplified with primers KMp207 and KMp181. The ORF and promoter segments were amplified along with 40bp of complementary overlapping sequence. Products from the ORF and promoter reactions were mixed 1:1 and amplified using primers KMp178 (*E. coli*) or KMp177 (*S*. Typhimurium) and KMp181 to amplify the promoter/ORF chimeras. The resulting fragment and pTH19Kr (69) were digested with *Bam*HI and *Hin*dIII and ligated together, generating pKM05 (*E. coli* promoter) and pKM07 (*S.* Typhimurium promoter). Plasmids were transformed into *S*. Typhimurium 14028s *slyA* for gene expression analysis. The *hnsT* coding sequence was amplified from E2348/69 genomic DNA using the oligonucleotides EcoRI-hnsT-F and HindIII-hnsT-R. Both pBAD18 and the resulting PCR fragment were digested with *Eco*RI and *Hin*dIII, agarose gel-purified, and ligated together to generate pRW57.

### Fusaric acid resistance assays

Cultures were grown overnight in LB broth at 37°C, then diluted 1:100. Thirty microliters of this dilution were added to 270 µl of LB containing 30 µg/ml freshly prepared fusaric acid in 100-well BioScreen plates (GrowthCurves, Piscataway, NJ). Cultures were grown with continuous maximum shaking at 37°C and regular OD_600_ measurements were taken on a BioScreen C MBR. Fresh 20 mg/ml stock solutions of fusaric acid were prepared in DMF (dimethyl formamide).

### NMR analysis of SlyA-salicylate

SlyA protein for NMR analysis was prepared as an *N*-terminal 6×His-tagged protein from cells grown in M9 minimal medium supplemented with ^15^N-ammonium chloride (Cambridge Isotope Labs, Tewksbury MA). SlyA protein expression was induced by the addition of 2 mM IPTG (isopropyl-ß-D-thiogalactopyranoside) and the protein purified to homogeneity using Ni-affinity chromatography as previously described (21), except that samples were dialyzed in 50 mM Tris-HCl, pH 8.0, 150 mM NaCl, 0.1mM EDTA, and 3 mM DTT following purification. Protein and ligand solutions for NMR experiments were prepared in 25 mM sodium phosphate, 150 mM NaCl buffer at pH 7.0 containing 10% D_2_O. NMR spectra were collected on a Bruker DMX 500hHz spectrometer (Bruker, Billerica MA) on samples equilibrated at 35°C and consisting of 250-350 µM ^15^N-labeled His-SlyA in the absence or presence of 2mM sodium salicylate or 0.5mM sodium benzoate. Spectra were processed using NMR-Pipe (70) and analyzed using NMR-View (71).

### SlyA-salicylate crystallization

SlyA was over-expressed and purified for crystallization as previously described (21), except that samples were dialyzed in 50 mM Tris-HCl, pH 8.0, 150 mM NaCl, 0.1 mM EDTA, and 3 mM DTT following purification. Cryo I and II sparse matrix crystallization screens (Emerald Biosystems, Bainbridge Island, WA) were used to determine initial conditions for His-SlyA crystal formation by sitting-drop vapor diffusion in 24-well crystal trays. Equal volumes (4µL) of SlyA and crystallization solutions were mixed before plates were sealed and kept at room temperature. Crystals appeared within 4-7 days in 20% PEG 300, 10% glycerol, 0.1 M phosphate/citrate buffer, pH 4.2, 0.2M ammonium sulfate (condition #14). Subsequent crystal growth was performed using lab-made 20% PEG 400, 10% glycerol, 0.1 M phosphate/citrate buffer, pH 4.2, and 0.2 M ammonium sulfate. Crystallization of SlyA with sodium salicylate was achieved by adding dilutions of a 2M sodium salicylate aqueous stock solution to the protein-crystallization solution mixture. SlyA-salicylate crystals appeared within 7-10 days. A single crystal appeared at a concentration of 75 mM sodium salicylate and grew to a maximum size of 500µ × 300µ × 150µ (Fig. S4). This crystal was used for X-ray diffraction experiments.

The crystal was frozen at 100°K in its crystallization solution for diffraction data collection on GM/CA-CAT beamline 23-ID-D at the Advanced Photon Source. The space group for the crystals is P2_1_2_1_2 with two SlyA molecules in the asymmetric unit. The diffraction data were processed with HKL2000 (72). Dataset statistics are shown in Table S1. The crystal structure of the salicylate complex of SlyA was solved using a model of a previously investigated structure of apo-SlyA (unpublished data). That apo-structure was solved using the molecular replacement program, MOLREP (73) with a search model generated by applying Swiss-Model (74) and the SlyA amino acid sequence to PDB entry 2FBH (75). The structural model for the salicylate complex was refined using REFMAC-5 (76) in the CCP4 suite (77). Rfree (78) was calculated using 5% of the data in the test set. A high-resolution limit of 2.3Å was applied for the refinement, consistent with standards appropriate when the structure was solved. This is the resolution at which Rmerge for the data set drops below 0.40. XtalView (79) and Coot (80) were used to examine sigma A weighted |Fo|-|Fc| and 2|Fo|-|Fc| electron density maps (81). MOLSCRIPT(82), and Raster3D (83) were used to produce structural figures for this paper. Table S2 contains refinement statistics for the structure. Coordinates and structure factors have been deposited in the Protein Data Bank with identifier 3DEU.

### 6×His-RovA purification

RovA was purified using the same protocol as described previously for SlyA (17). However, overexpression cultures were grown at 24°C to OD_600_=0.5. IPTG was added to a final concentration of 1mM, and cultures were incubated at 24°C for an additional four h before the cells were harvested by centrifugation and cell pellets stored at −80°C for the subsequent purification of RovA.

### *In vitro* transcription

IVT assays were performed essentially as previously described (17) with the following modifications. All SlyA IVT assays were performed at 37°C. All RovA IVT assays were performed at 24°C. Where indicated, sodium salicylate was added to IVT reactions prior to the addition of SlyA or RovA. All oligonucleotides and templates used in IVT reactions are indicated in Table S6.

### RNA isolation and qRT-PCR

RNA was purified using Trizol (Life Technologies, Carlsbad CA) according to the manufacturer’s protocols. cDNA was generated using the QuantiTect Reverse Transcription Kit (Qiagen, Hilden, Germany), and quantified in a BioRad CFX96 (BioRad, Hercules CA), using SYBR Green Master Mix (84).

For the analysis of *pagC* expression under inducing conditions, *S*. Typhimurium cultures were grown to early stationary phase (OD_600_≈2.0) at 37°C in LB broth, then washed three times in N-minimal medium containing either 10 µM (inducing) or 10 mM MgCl_2_ (non-inducing). Cultures were re-suspended in the appropriate N-minimal medium with 2mM sodium salicylate where indicated and incubated for an additional 30 min at 37°C before cells were harvested for RNA purification studies. For the complementation of *slyA* with *slyA* or *rovA in trans*, overnight cultures were diluted to 0.05 OD_600_ and grown for two h at 24°C before adding arabinose to a final concentration of 0.02% w/v. Cultures were grown for an additional six h before harvesting RNA.

*Y. pseudotuberculosis* strains YPIII and YP107 were diluted from overnight cultures to 0.05 OD_600_ in LB and grown with shaking at 24°C. For the inhibition of H-NS via H-NST_EPEC_ over-expression, and complementation of *rovA* with *rovA* or *slyA in trans*, YPIII and YP107 cultures carrying either pBAD18 or pRW57 were grown for two hrs before arabinose was added to a final concentration of 0.02% w/v. Cultures were grown for an additional six hrs before harvesting RNA. For *slyA*/*rovA* complementation studies, rpoAYS primers targeting *rpoA* were used as loading controls, as *rpoA* is sufficiently conserved between the two species as to allow use of the same primers.

## Supporting information

## Acknowledgments

The National Institutes of Health provided support to FCF (AI39557, AI44486, AI118962, AI112640) and SJL (A148622). GM/CA@APS has been funded in whole or in part with Federal funds from the National Cancer Institute (ACB-12002) and the National Institute of General Medical Sciences (AGM-12006). This research used resources of the Advanced Photon Source, a U.S. Department of Energy (DOE) Office of Science User Facility operated for the DOE Office of Science by Argonne National Laboratory under Contract No. DE-AC02-06CH11357. We thank Eric Larson for helpful discussions and suggestions. We also thank Catherine Eakin and Ponni Rajagopal for assistance with protein purification and crystal growth, Petra Dersch for kindly providing *Y. pseudotuberculosis* YPIII and YP107, and Ralph Isberg for providing *Y. pseudotuberculosis* IP32953. Finally, we thank Dr. Rachel Klevit for use of her NMR facilities to collect spectra of SlyA-salicylate complexes.

## Notes

#### Summary of Updates

Signifcant additional data (Figure 2 - figure supplement 2, Figures 4-6) and discussion. Authors updated.

