## Supplementary material for "The Evolution of SlyA/RovA Transcription Factors From Repressors to Counter-Silencers in *Enterobacteriaceae*"

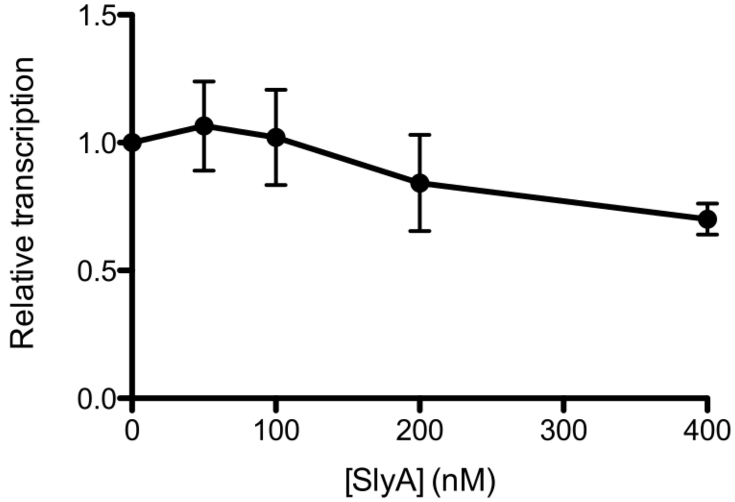


**Figure 1 – figure supplement 1. SlyA does not activate *pagC* transcription.** To confirm that SlyA does not function as an activator at up-regulated targets, *in vitro* transcription (IVT) assays of the model *S*. Typhimurium SlyA-regulated *pagC* promoter were performed in the presence of increasing SlyA concentrations. Reactions were assembled as previously described (1) using the *pagC* target plasmid, pRW6. Data are normalized to the RNAP control reaction and represent the mean ± SEM; n=3.

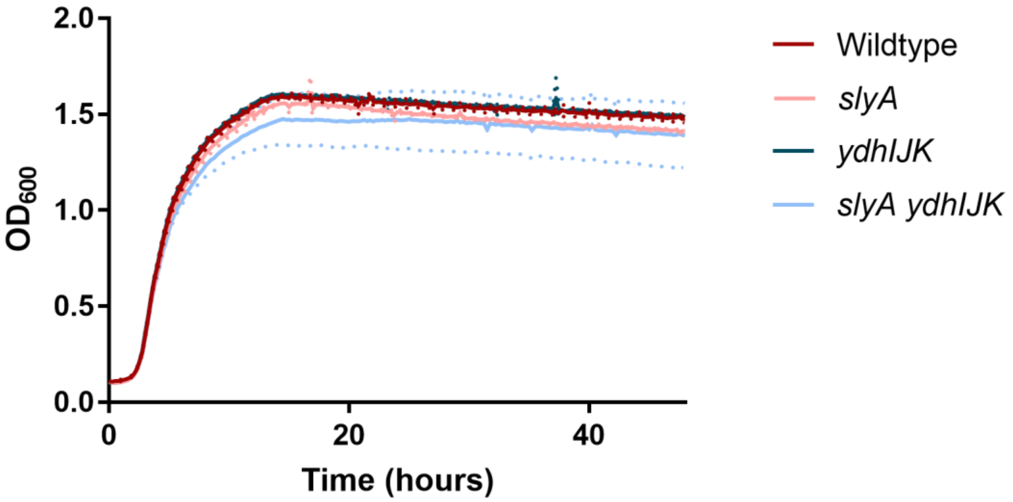


**Figure 1 – figure supplement 2. Mutations of *slyA* or *ydhIJK* do not significantly affect *S.* Typhimurium growth.** Culture densities ($OD_{600}$) of wildtype, *slyA*, *ydhIJK*, and *slyA ydhIJK* cultures grown at 37°C were monitored to determine whether the mutation of either *slyA* or *ydhIJK* affects bacterial growth. Solid lines represent the mean of three independent experiments, each consisting of three technical replicates. Dashed lines represent the SD.

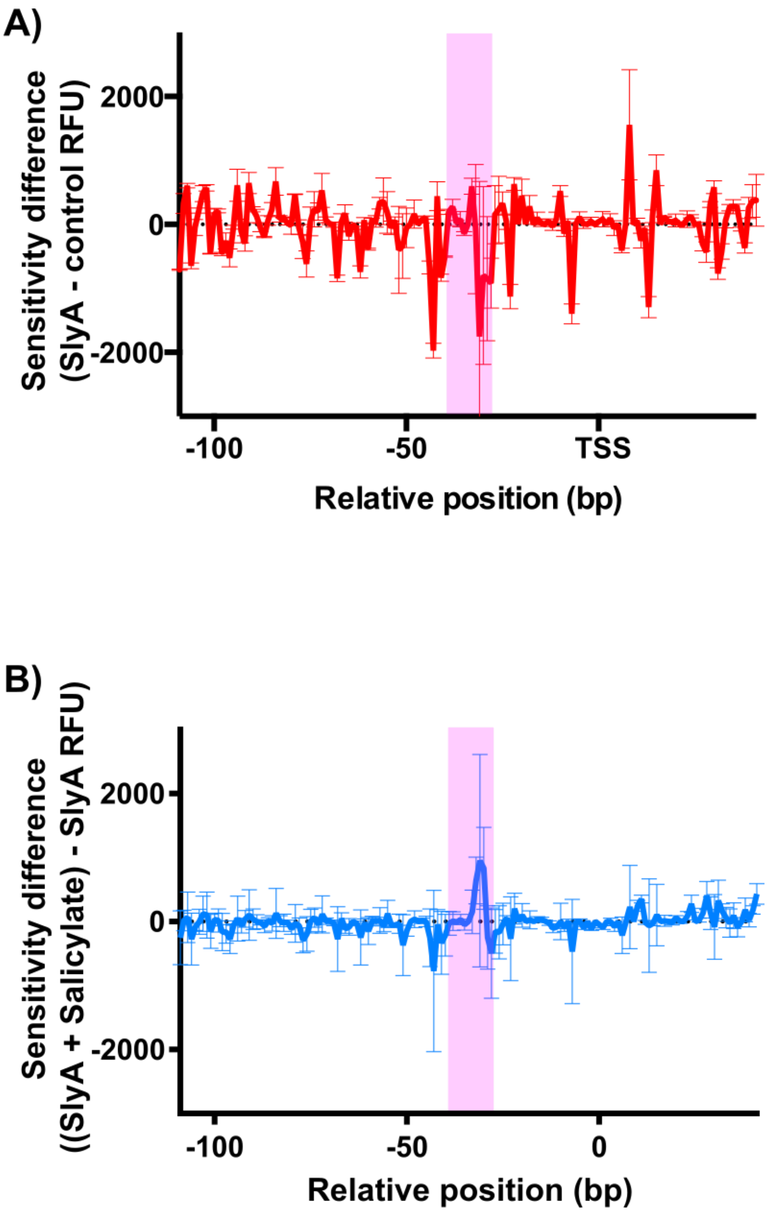


**Figure 2 – figure supplement 1. Salicylate destabilizes the SlyA-DNA complex.** *In vitro* DNase I footprinting was performed on the *ydhI* promoter region in the presence of 2 µM SlyA (A). Results are presented as a Differential DNA Footprint Analysis (DDFA) plot, representing the difference in fluorescence in RFU (relative fluorescence units) between SlyA-bound DNA and an unbound DNA control. Values on the horizontal axis indicate the relative distance from the *ydhI* transcriptional start site (TSS). Downward peaks indicate sites of protection from DNase I digestion, and upward peaks indicate regions of increased DNase I sensitivity, suggesting bending or distortion of the DNA. To determine how salicylate influences the SlyA-DNA interaction, DNase I footprinting was also performed on the SlyA-DNA complex in the presence of 2 mM sodium salicylate (B). Results are presented as a DDFA plot representing the difference in RFU between SlyA-bound DNA in the presence or absence of salicylate. There are few significant differences except for a destabilization of the SlyA-DNA interaction between bases -

30 to -32 in the presence of salicylate. This indicates that SlyA is still able to interact with DNA in the presence of salicylate, but SlyA binding near positions -30 to -32 is destabilized and likely to be critical for SlyA-mediated repression of *ydhI*. Sequence analysis reveals a region with 75% homology to the high affinity consensus SlyA binding site (5’- TTAGCAAGCTAA-3’) between bases -28 and -40 (5’- TTGGTAAGCAAA-3’) (2-4). This region is highlighted in pink in each panel. Data represent the mean ± SD; n=3 independent replicates.

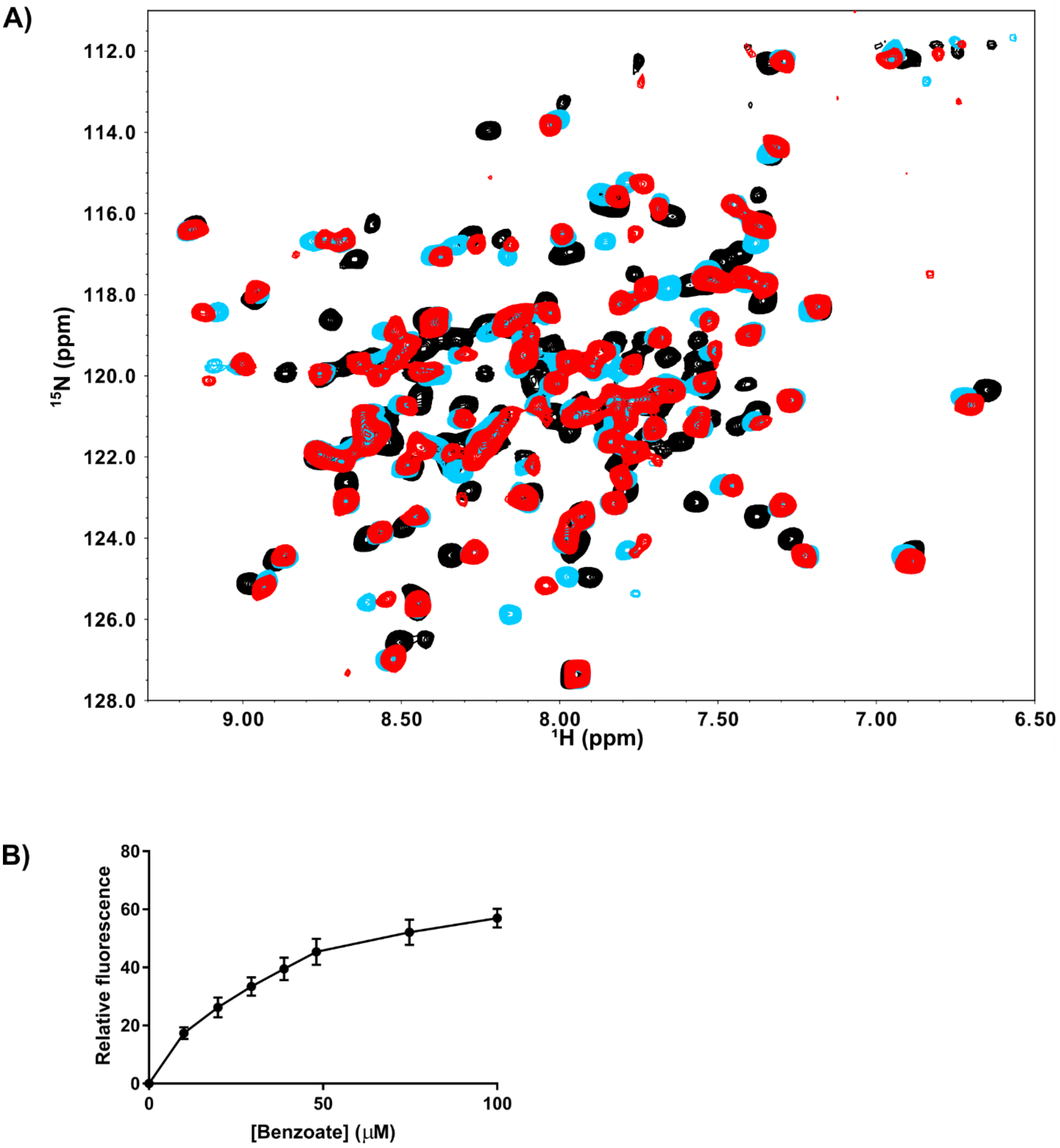


**Figure 2 – figure supplement 2. The affinity of the SlyA-ligand interaction.** The benzoate-SlyA interaction was examined to determine the affinity of the SlyA ligand interaction. To confirm that benzoate induces similar conformational changes in SlyA to those observed with salicylate, $^{1}H,^{15}N$-TROSY NMR spectroscopy was performed on 0.3mM uniformly-labelled $^{15}N$-SlyA in the absence (black) or presence (blue) of 0.5mM benzoate (A). SlyA in the presence of 2mM salicylate (red) is overlaid for comparison. To determine the affinity of the SlyA-benzoate interaction, the change in maximum intensity of intrinsic tryptophan fluorescence ($l_{ex}$ = 290nm) of SlyA was measured in relation to increasing concentrations of benzoate (B). The data were fit to a standard hyperbolic binding isotherm yielding a $K_D$ of ~40μM. Data represent the mean ± SD; n=2.

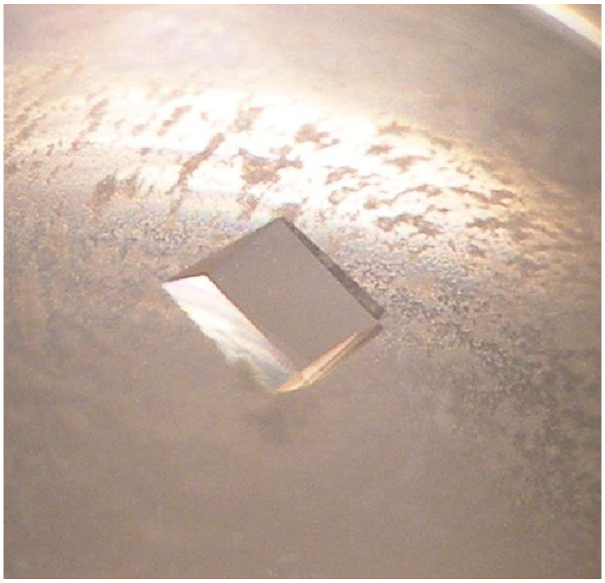

**Figure 3 – figure supplement 1. Crystallization of SlyA-salicylate.** SlyA-salicylate was crystalized at room temperature in 20% PEG 400, 10% glycerol, 0.1M phosphate/citrate buffer, pH 4.2, 0.2M ammonium sulfate, and 75 mM sodium salicylate. This crystal was used to determine the structure of SlyA-salicylate.

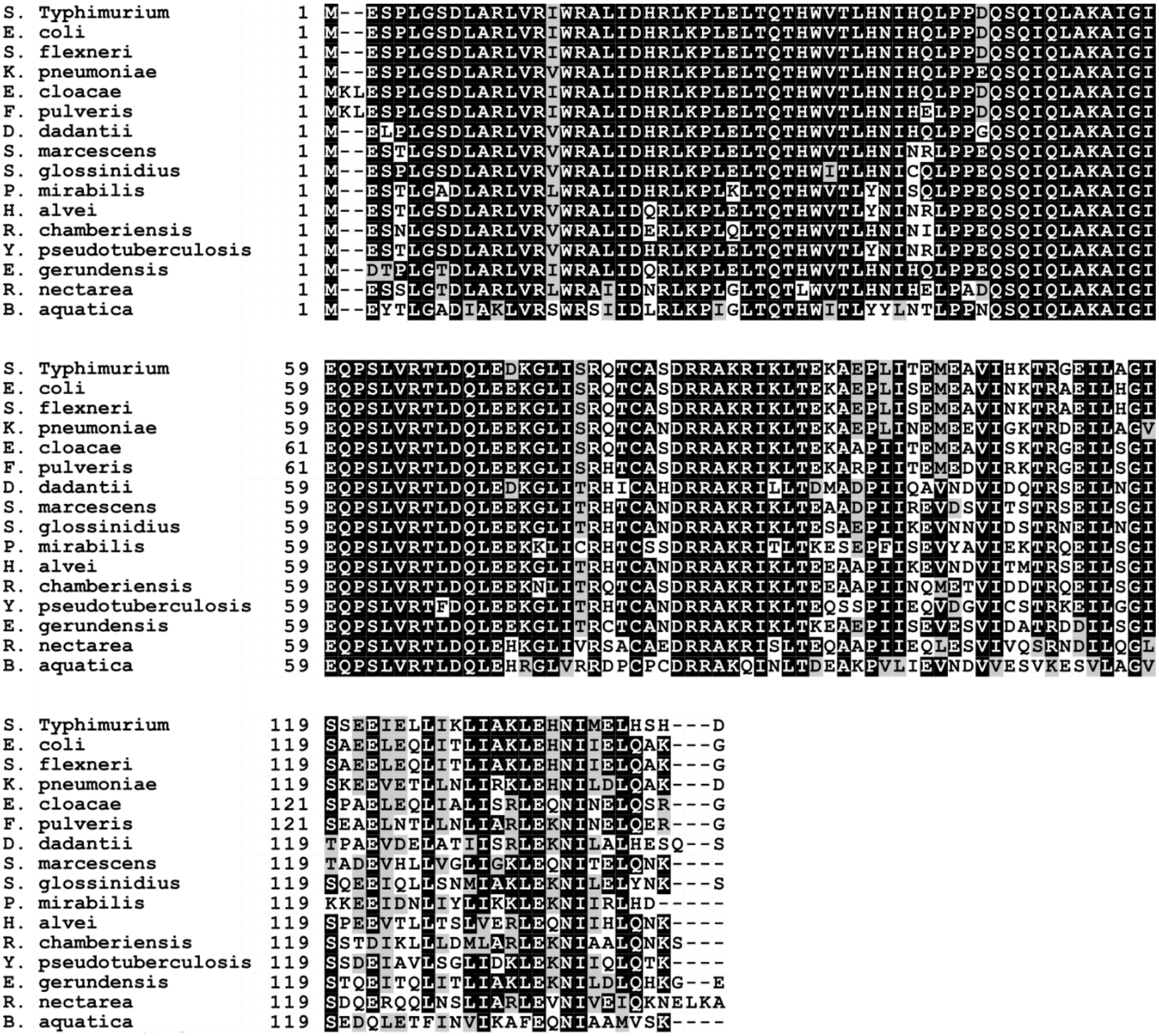


**Figure 5 – figure supplement 1. Multiple alignment of SlyA orthologs in *Enterobacteriaceae.*** The protein sequences of SlyA orthologs from representative species of *Enterobacteriaceae* were aligned using T-Coffee (5). Amino acid positions are indicated to the left of each sequence.

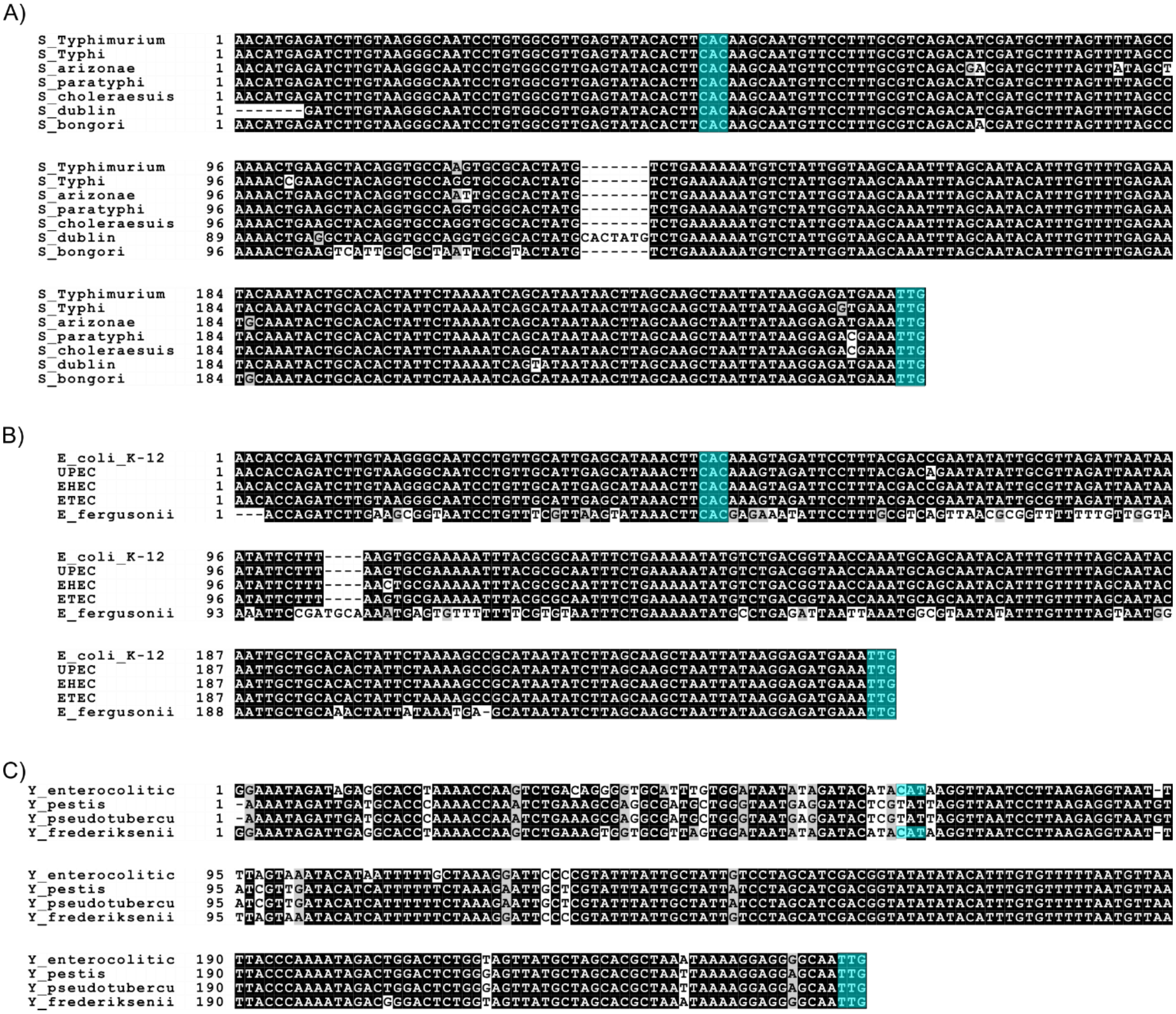


**Figure 6 – figure supplement 1. Multiple alignment of the *slyA* regulatory regions of *Enterobacteriaceae*.** The 250 bp immediately upstream of the *slyA* ortholog coding sequence from representative species of A) *Salmonella*, B) *Escherichia*, and *Yersinia* were aligned using Pro-Coffee (5). The start codons of *slyA* (positions 248-250) and *ydhI* (species dependent; positions 47-70 when present) are highlighted in teal.

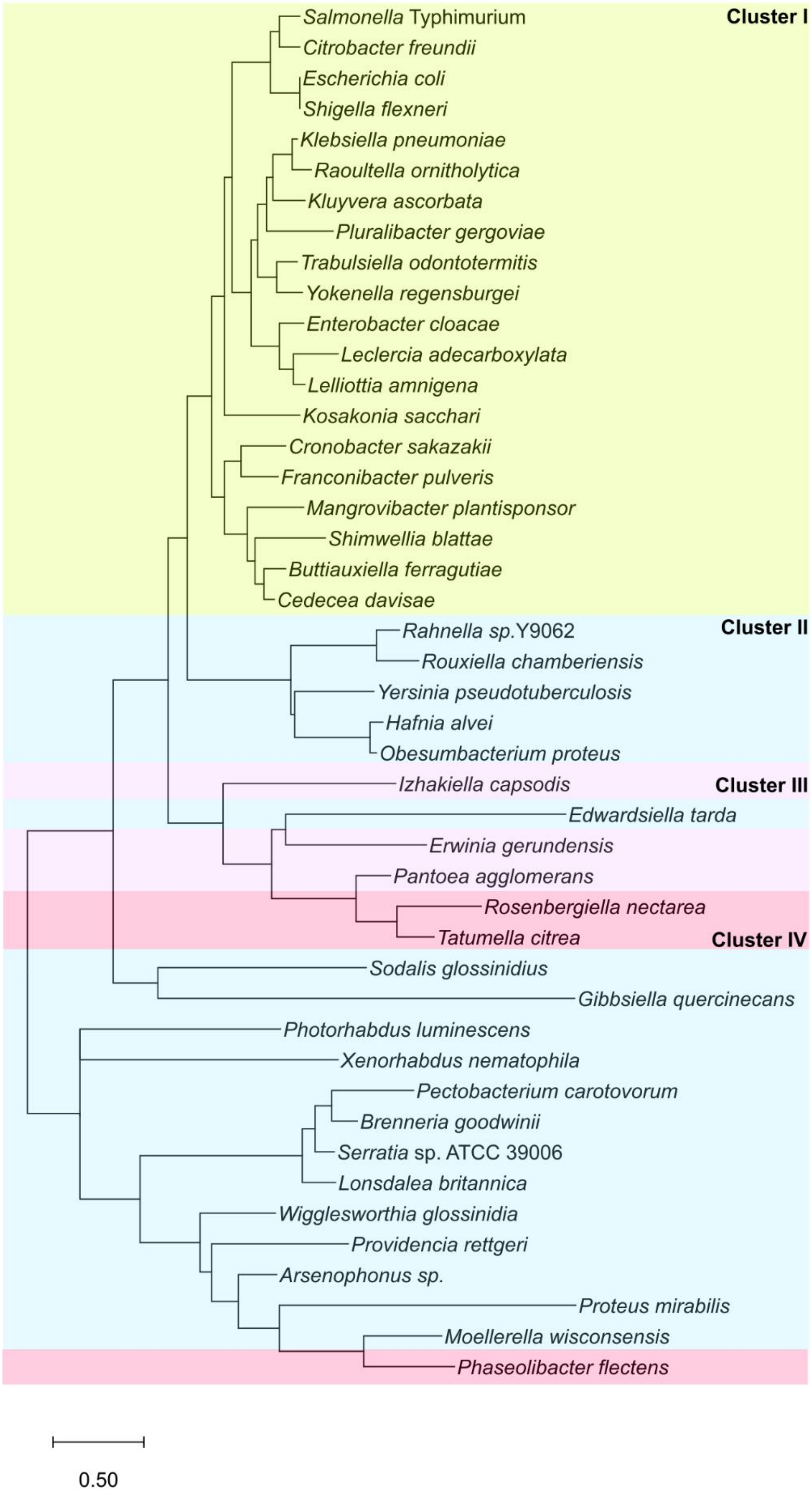
Cluster I
Salmonella Typhimurium
Citrobacter freundii
Escherichia coli
Shigella flexneri
Klebsiella pneumoniae
Raoultella ornitholytica
Kluyvera ascorbata
Pluralibacter gergoviae
Trabulsiella odontotermitis
Yokenella regensburgei
Enterobacter cloacae
Leclercia adecarboxylata
Lelliottia amnigena
Kosakonia sacchari
Cronobacter sakazakii
Franconibacter pulveris
Mangrovibacter plantisponsor
Shimwellia blattae
Buttiauxiella ferragutiae
Cedecea davisae
Cluster II
Rahnella sp.Y9062
Rouxiella chamberiensis
Yersinia pseudotuberculosis
Hafnia alvei
Obesumbacterium proteus
Izhakiella capsodis
Cluster III
Edwardsiella tarda
Erwinia gerundensis
Pantoea agglomerans
Rosenbergiella nectarea
Tatumella citrea
Cluster IV
Sodalis glossinidius
Gibbsiella quercinecans
Photorhabdus luminescens
Xenorhabdus nematophila
Pectobacterium carotovorum
Brenneria goodwinii
Serratia sp. ATCC 39006
Lonsdalea britannica
Wigglesworthia glossinidia
Providencia rettgeri
Arsenophonus sp.
Proteus mirabilis
Moellerella wisconsensis
Phaseolibacter flectens
0.50

91

**Figure 6 – figure supplement 2. Phylogenetic analyses of the *slyA* promoters of *Enterobacteriaceae*.** Species from genera analyzed in Figure 5 for which annotated genomic data were available on BioCyc (6) were subsequently used in phylogenetic analyses of the 250 bp region immediately upstream of the *slyA* coding sequence, containing the *slyA* promoter. The evolutionary history of the *slyA* promoter region was inferred using the Maximum Likelihood method and the Tamura-Nei model (7) with the MEGA X software (8). The tree is drawn to scale with branch lengths measured in the number of substitutions per site. Branches are highlighted to indicate the corresponding SlyA cluster (Figure 5). The promoter regions of some members of clusters III and IV are more closely related to members of cluster II than to other members of their own cluster, suggesting that similar promoters may have evolved in parallel in distantly-related species.

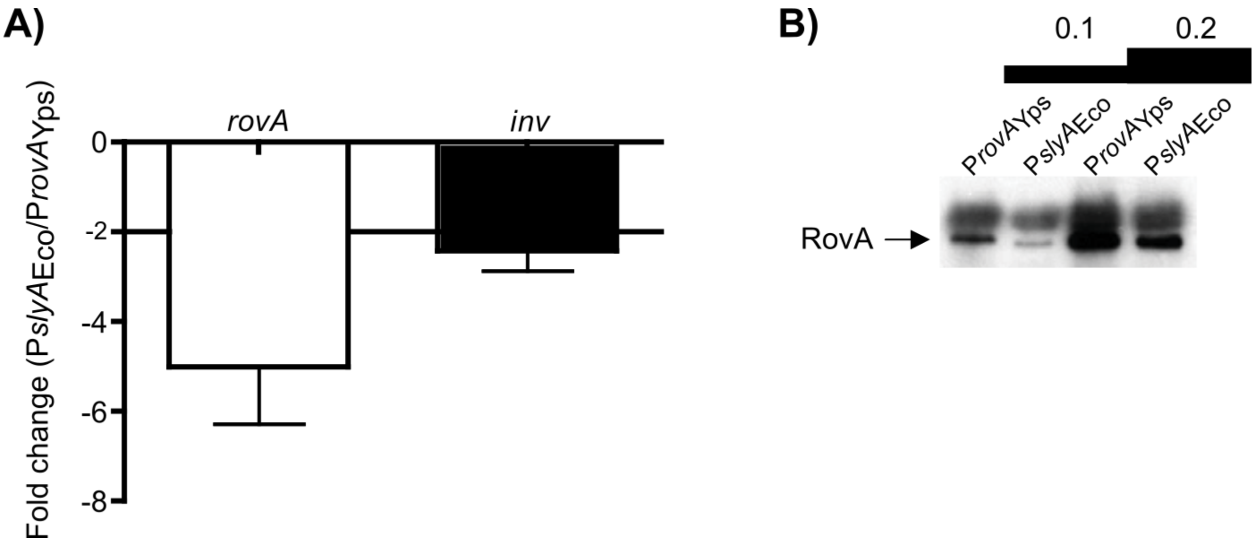


**Figure 8 – figure supplement 1. The endogenous *rovA* promoter of *Y. pseudotuberculosis* expresses *rovA* more strongly than the *E. coli slyA* promoter.** To compare the strength of the native *rovA* promoter (P*rovA*$_{Yps}$) and P*slyA*$_{Eco}$, the native *rovA* promoter was replaced with the corresponding sequence from *E. coli* and the resulting chimera examined for its ability to regulate the RovA targets *rovA* and *inv* in *Y. pseudotuberculosis*. mRNA was purified from cultures grown for 6 h at 26°C in LB, and transcripts were quantified by qRT-PCR and normalized to *gyrB* (A). RovA protein levels were confirmed by immunoblot analysis using 0.1 and 0.2 OD$_{600}$ of culture after 6 h of growth (left two and right two lanes, respectively) (B). The position of RovA is indicated by an arrow.

**Table S1. Data Collection Statistics.**

| | |
|---|---|
| Space group | $P2_12_12$ |
| Unit cell dimensions (a,b,c) (Å) | 63.38 78.02 84.39 |
| Molecules per asymmetric unit | 2 |
| Resolution (Å) (last shell) | 50.0-2.00 Å (2.07-2.00) |
| Unique reflections (last shell) | 27998 (2583) |
| Completeness (last shell) | 96.4% (90.8) |
| Redundancy (last shell) | 12.9 (7.1) |
| $\langle I \rangle / \langle \sigma(I) \rangle$ (last shell) | 31.7 (1.1) |
| $R_{merge}$ (last shell) | 0.074 (>1.00) |

**Table S2. Refinement Statistics.**

| | |
|---|---|
| Resolution | 20.0 – 2.3 |
| R factor (working set) | 0.228 |
| Rfree (test set=5% of the overall) | 0.259 |
| # unique reflections | 17811 |
| Number of protein atoms | 2110 |
| Number of solvent atoms | 19 |
| Number of heteroatoms | 60 (6 salicylates) |
| Wilson B value | 51.8 $Å^2$ |
| Average B value from refinement | 64.7 $Å^2$ |
| Ramachandran quality | 97.4% in most-favored regions<br>2.2% in additional allowed regions<br>0.4% in generously allowed regions |
| rms deviation - bond lengths | 0.011 Å |
| rms deviation - bond angles | 2.0 ° |

**Table S3. Polymorphisms in salicylate binding sites I and II of SlyA lineage TFs.**

| Site I | T32 | V35 | T36 | I56 | I58 | S62 | T66 |
|---|---|---|---|---|---|---|---|
| *Budvicia aquatica* | I | | | | | | |
| *Gibbsiella quercinecans* | I | | | | | | |
| *Phaseolibacter flectens* | I | | | | | | |
| *Pragia fontium* | I | | | | | | |
| *Sodalis glossinidius* | I | | | | | | |
| *Wigglesworthia glossinidia* | I | | | | | | |

| Site II | G6’ | S7’ | A10’ | R14’ | R17’ | W16 | I20 | W34 | V35 | H38 |
|---|---|---|---|---|---|---|---|---|---|---|
| *Arsenophonus sp.* | | T | | | | | | | | Y |
| *Budvicia aquatica* | | A | | | | | | | I | Y |
| *Cosenzaea myxofaciens* | | A | | | | | | | | Y |
| *Cronobacter sakazakii* | | | T | | | | | | | |
| *Edwardsiella tarda* | | | S | | | | | | | Y |
| *Enterobacillus trilobii* | | | | | | | | | | Y |
| *Erwinia gerundensis* | | T | | | | | | | | |
| *Franconia pulveris* | | | | | | | | | I | |
| *Hafnia alvei* | | | | | | | | | | Y |
| *Izhakiella australiensis* | | T | S | | | | | | | |
| *Leminorella grimontii* | | A | | | | | | | | Y |
| *Moellerella wisconsensis* | | T | | | | | | | | |
| *Obesumbacterium proteus* | | | | | | | | | | Y |
| *Pantoea agglomerans* | | T | S | | | | | | I | |
| *Pectobacterium carotovorum* | | | | | | | V | | | |
| *Phaseolibacter flectens* | S | I | | | | | | | | |
| *Photorhabdus luminescens* | | | | | | | | | | Y |
| *Pragia fontium* | | A | | | | | | | I | |
| *Proteus mirabilis* | | A | | | | | | | | Y |
| *Providencia rettgeri* | | T | S | | | | | | | Y |
| *Rosenbergiella nectarea* | | T | | | | | | | | |
| *Shimwellia blattae* | | | S | | | | | | | |
| *Sodalis glossinidius* | | | | | | | | | I | |
| *Tatumella ptyseos* | | T | | | | | | | | |
| *Trabulsiella guamensis* | | | | | | | | | I | |
| *Wigglesworthia glossinidia* | | | S | | | | | | | |
| *Xenorhabdus ishibashii* | | | | | | | | | | Y |
| *Yersinia pseudotuberculosis* | | | | | | | | | | Y |

**Table S4. Oligonucleotides used in this study.**

| Name | Sequence (5’-3’) |
|---|---|
| 6-FAM-ydhI-R | 56-FAM/ATTGAGCAGCCATCGCTGG |
| 6HisRovA-F | CTACCATGGAACATCACCATCACCATCACGGTGCTGGAGCATTGGAATCGACATTAGG |
| 6HisRovA-R | CTAGGATCCTTACTTAGTTTGTAATTGAATAATATTTTTC |
| A10P-F | CTAGGTTCTGATCTGCCGCGGTTGGTGCGCATTTGG |
| A10P-R | CCAAATGCGCACCAACCGCGGCAGATCAGAACCTAG |
| BamHI-inv-F | TGCGGATCCTTGGGTAGCGGATAATATTG |
| BamHI-rovA-F | TGCGGATCCGTTGCTGACGAGATG |
| C81A-F | GGCTAATTTCGCGGCAAACCGGCGCCAGCGATCGTCGCGCTAAG |
| C81A-R | CTTAGCGCGACGATCGCTGGCGGAGGTTTGCCGCGAAATTAGCC |
| EcoRI-hnsT-F | AGTCGAATTCAAGGAGCAAAAAAATGATTGATGAATTTC |
| EcoRI-inv-R | TGCAGAATTCGCCGTTGCCCCTCC |
| EcoRI-rovA-F | TACGGAATTCAAGGAGGAGCAATTGGAATCGACATTAGGATCTG |
| EcoRI-rovA-R | TGCAGAATTCGCCATTGGAACAATCTTG |
| EcoRI-slyA-F | TACGGAATTCAAGGAGGAGCAATTGGAATCGCCACTAGGTTCTG |
| G6A-F | GAAATTGGAATCGCCACTAGCTTCTGATCTGGCACGGTTGG |
| G6A-R | CCAACCGTGCCAGATCAGAAGCTAGTGGCGATTCCAATTTC |
| H38A-F | ACACATTGGGTCACGTTGGCGAATATTCATCAATTGCCG |
| H38A-R | CGGCAATTGATGAATATTCGCCAACGTGACCCAATGTGT |
| HindIII-hnsT-R | GCATAAGCTTCAGTCAATGAGATCTTCTGGCG |
| JKP736 | GACCACACCCGTCCTGTGTGAATGGCTATCTACCAGGG |
| JKP737 | CGCTGGCGCTGGTTTGCCGCGAAATTAGCC |
| JKP738 | CGGCAAACCAGCGCCAGCGATCGTCGCG |
| JKP739 | GATGCGTCCGGCGTAGAGGCCGGCCTAACTGGGTAT |
| KMp177 | CGGATCCCCAGATAACGAACCCAAGCG |
| KMp178 | CGGTCCCCAGATGACGAATCCAAACG |
| KMp181 | CAAGCTTTGGTCACATGGCCACACGTAT |
| KMp206 | TGGCGATTCCAATTTCATCTCCTTATAATTAGCTTGCTAAG |
| KMp207 | CTTAGCAAGCTAATTATAAGGAGATGAAATTGGAATCGCCA |
| KpnI-rovA-R | GAGGTACCTTACTTAGTTTGTAATTGAATAATATTTTTCTC |
| KpnI-slyA-R | GAGGTACCTCAATCGTGAGAGTGCAATTCCATAATATTGTGTTC |
| pagC-3'-F | AGAACATTCCACTCAGGATGGCGA |
| pagC-3'-P | TGTAGAGGAGATGTTGCTTCC |
| pagC-3'-R | GACGACGATATTCTCCAGCGGATT |
| rpoAYS-F | CCAAGGTGACCCTTGAGCC |
| rpoAYS-R | TAGTACACCATCAATCTCAACCTCGG |
| R14A-F | CTGGCACGGTTGGTGCAATTGGCGTGCTCTG |
| R14A-R | CAGAGCACGCCAAATTGCCACCAACCGTGCCAG |
| slyA-F | TTGGCTGGGATTTCTTCAGAG |
| slyA-R | TCGTGAGAGTGCAATTCCATAA |
| slyAcomp-F | GGTGCCAAGTGCGCACTATCTCTG |
| slyAcomp-R | GTATGCCCTGCACCTCAATCGTG |
| S7A-F | GGCATTGAGCAGCCAGCGCTGGTACGCACGTTGGATC |
| S7A-R | GATCCAACGTGCGTACCAGCGCTGGCTGCTCAATGCC |

| | |
|---|---|
| SlyAreg-F | CGCGGGATCCTTACCGCTGTCCAATGGCTAC |
| SlyAreg-R | CGCGAAGCTTTTCCGACTTCGTTTAAGATTGG |
| STM-Eco-slyA-F | CAGCATAATAACTTAGCAAGCTAATTATAAGGAGATGAAATTGGAATCGCCACTAGGTTC |
| STM-Eco-slyA-R | CTTTACGTGTGGTCACATGGCCACACGTATGCCCCTGCACCTCACCCTTTGGCCTGTAA |
| STM-slyA-targ-F | ATCAGCATAATAACTTAGCAAGCTAATTATAAGGAGATGAAATAGACAGCTGCATGCAT |
| STM-slyA-targ-R | CTTTACGTGTGGTCACATGGCCACACGTATGCCCCTGCACCTCAGTGTAGGCTGGAGCTG |
| T66A-F | CAGCCATCGCTGGTACGCGCGTTGGATCAACTTGAAGATAAG |
| T66A-R | CTTATCTTCAAGTTGATCCAACGCGCGTACCAGCGATGGCTG |
| W34A-F | TTGACGCAGACACATGCGGTCACGTTGCACAATATTC |
| W34A-R | GAATATTGTGCAACGTGACCGCATGTGTCTGCGTCAA |
| WNp318 | TTGTGAAGTGTATACTCAACGCCACAGGATTGCCCTTACACATATGAATATCCTCCTTAG |
| WNp319 | GTCGCAGATACGCTGTAGTTCCTGTAGCGTGACGGCAAGCGTGTAGGCTGGAGCTGCTTC |
| ydhI-F | GTAAAGAGGGAGAGATCCATTAACA |
| ydhI-R | TGCGCTTGGGTTCGTTAT |

**Table S5. Plasmids used in this study.**

| Name | Description | Source |
|---|---|---|
| pBAD18 | Arabinose-inducible vector | (9) |
| pET16b::*slyA* | pET16b::6×His-*slyA* | (10) |
| pJK723 | Suicide construct containing C81S mutation | This study |
| pKM05 | 14028s $P_{slyA}$-14028s *slyA* ORF chimera | This study |
| pKM07 | *E. coli* K-12 $P_{slyA}$-14028s *slyA* ORF chimera | This study |
| pRDH10 | Suicide vector | (11) |
| pRW6 | *pagC* IVT target | (1) |
| pRW20 | IVT scaffold vector | (1) |
| pRW39 | *slyA*/*ydhI* region IVT target | This study |
| pRW54 | *rovA* region IVT target | This study |
| pRW55 | *inv* region IVT target | This study |
| pRW57 | pBAD18::$hnsT_{EPEC}$ | This study |
| pRW58 | pBAD18::*slyA* | This study |
| pRW59 | pBAD18::*rovA* | This study |
| pRW60 | pTRC99::6×His-*rovA* | This study |
| pSL2143 | pWSK29 *slyA* | This study |
| pSL2143-G6A | pWSK29 *slyA* G6A | This study |
| pSL2143-S7A | pWSK29 *slyA* S7A | This study |
| pSL2143-A10P | pWSK29 *slyA* A10P | This study |
| pSL2143-R14A | pWSK29 *slyA* R14A | This study |
| pSL2143-H38A | pWSK29 *slyA* H38A | This study |
| pSL2143-T66A | pWSK29 *slyA* T66A | This study |
| pSL2143-C81A | pWSK29 *slyA* C81A | This study |
| pSR47s | Suicide vector | (12) |
| pSW172 | Temperature sensitive plasmid | (13) |
| pTH19Kr | Low copy number vector | (14) |
| pTRC99a | Tightly controlled expression vector | (15) |
| pWSK29 | Low copy number vector | (16) |

**Table S6. Components of IVT reactions.**

| Target promoter | Template | Probe | Primers |
|---|---|---|---|
| *pagC* | pRW6 | pagC-3’-P | pagC-3’-F/pagC-3’-R |
| *slyA* | pRW39 | slyA-R | slyA-F/slyA-R |
| *ydhI* | pRW39 | ydhI-R | ydhI-F/ydhI-R |
| *inv* | pRW55 | inv-R | inv-F/inv-R |
| *rovA* | pRW54 | rovA-R | rovA-F/rovA-R |

**Supplementary Methods.**

**DNase I footprinting and DDFA.** DNase I footprinting was performed using methods described previously (1). SlyA was incubated at a final concentration of 2 µM in the presence or absence of sodium salicylate with the target plasmid pRW39 for 10 min at 37°C. DNase I was added and allowed to digest the SlyA-DNA complex for two min at 37°C before the reaction was quenched with cold stop buffer, purified, amplified by fluorescent primer extension using oligonucleotide 6-FAM-ydhI-R and analyzed by DDFA.

**Ligand Binding Assays**. To estimate the $K_D$ of ligand binding to SlyA, the quenching of intrinsic tryptophan fluorescence in SlyA was measured in the presence of the aromatic carboxylate, benzoate. Benzoate was selected for these experiments because (1) $^{1}$H,$^{15}$N-TROSY NMR spectra show that binding of benzoate induces chemical shift perturbations similar to those observed with other aromatic carboxylates such as salicylate, and (2) inner filter effects during fluorescence measurements can be avoided since there is little overlap in the UV spectra of benzoate and SlyA, and an $l_{ex}$ of 290nm can be used to stimulate intrinsic SlyA tryptophan fluorescence. Titrations were conducted starting with 0.2µM His-SlyA in 25mM $NaPO_4$, 150mM NaCl, pH 7.0 at 25°C followed by sequential additions from an identical sample containing 2mM sodium benzoate. Emission spectra from 320 to 450nm ($l_{ex}$ = 290nm) were collected from samples ranging in benzoate concentration from 0-200mM, and the change in maximum intensity was plotted as a function of benzoate concentration. The data were fit to a standard hyperbolic binding isotherm yielding a $K_D$ of ~40µM.

***rovA* expression analysis.** *Y. pseudotuberculosis* IP32953 (17) was propagated at 26°C in LB. YPTB007 is a derivative of IP32953 in which the native *rovA* promoter was replaced with the *E. coli slyA* promoter. Briefly, the 691 bp 5' of the native *rovA* gene were replaced with the 292 bp 5' of the E. coli *slyA* gene using pSR47s (12) and homologous recombination. Replacement was confirmed by PCR amplification of the intergenic region and DNA sequencing.

For qRT-PCR and Western blot analyses, *Y. pseudotuberculosis* IP32953 and YPTB007 (n=3) were grown at 26°C with aeration. After 15 h, bacterial concentrations were determined by spectrophotometry and diluted to 0.1 $OD_{600}$ in fresh medium. Cultures were grown for 6 h at 26°C before cells were harvested for mRNA extraction. *rovA* and *inv* transcript levels were determined by qRT-PCR using SybrGreen and normalized to *gyrB* transcript levels (18, 19). For western blot analyses, 1.0 $OD_{600}$ equivalent of total bacteria was harvested and lysed in SDS protein loading buffer. 0.1 or 0.2 $OD_{600}$ equivalents were separated by SDS-PAGE, transferred to PVDF, and western blotting was performed with rabbit α-RovA antibody (1:1,000) (20).

**Supplementary References.**

13. Lopez CA, Winter SE, Rivera-Chávez F, Xavier MN, Poon V, Nuccio SP, Tsolis RM, Bäumler AJ. 2012. Phage-mediated acquisition of a type III secreted effector protein boosts growth of *Salmonella* by nitrate respiration. MBio 3.
14. Hashimoto-Gotoh T, Yamaguchi M, Yasojima K, Tsujimura A, Wakabayashi Y, Watanabe Y. 2000. A set of temperature sensitive-replication/-segregation and temperature resistant plasmid vectors with different copy numbers and in an isogenic background (chloramphenicol, kanamycin, *lacZ, repA*, *par*, *polA*). Gene 241:185-91.
15. Amann E, Ochs B, Abel KJ. 1988. Tightly regulated *tac* promoter vectors useful for the expression of unfused and fused proteins in *Escherichia coli*. Gene 69:301-15.
16. Wang RF, Kushner SR. 1991. Construction of versatile low-copy-number vectors for cloning, sequencing and gene expression in *Escherichia coli*. Gene 100:195-9.
17. Chain PS, Carniel E, Larimer FW, Lamerdin J, Stoutland PO, Regala WM, Georgescu AM, Vergez LM, Land ML, Motin VL, Brubaker RR, Fowler J, Hinnebusch J, Marceau M, Medigue C, Simonet M, Chenal-Francisque V, Souza B, Dacheux D, Elliott JM, Derbise A, Hauser LJ, Garcia E. 2004. Insights into the evolution of *Yersinia pestis* through whole-genome comparison with *Yersinia pseudotuberculosis*. Proc Natl Acad Sci U S A 101:13826-31.
18. Lawrenz MB, Lenz JD, Miller VL. 2009. A novel autotransporter adhesin is required for efficient colonization during bubonic plague. Infect Immun 77:317-26.
19. Lenz JD, Lawrenz MB, Cotter DG, Lane MC, Gonzalez RJ, Palacios M, Miller VL. 2011. Expression during host infection and localization of *Yersinia pestis* autotransporter proteins. J Bacteriol 193:5936-49.
20. Ellison DW, Miller VL. 2006. H-NS represses *inv* transcription in *Yersinia enterocolitica* through competition with RovA and interaction with YmoA. J Bacteriol 188:5101-12.
